# Smoother: A Unified and Modular Framework for Incorporating Structural Dependency in Spatial Omics Data

**DOI:** 10.1101/2022.10.25.513785

**Authors:** Jiayu Su, Jean-Baptiste Reynier, Xi Fu, Guojie Zhong, Jiahao Jiang, Rydberg Supo Escalante, Yiping Wang, Luis Aparicio, Benjamin Izar, David A Knowles, Raul Rabadan

## Abstract

Spatial omics technologies can help identify spatially organized biological processes, but existing computational approaches often overlook structural dependencies in the data. Here, we introduce Smoother, a unified framework that integrates positional information into non-spatial models via modular priors and losses. In simulated and real datasets, Smoother enables accurate data imputation, cell-type deconvolution, and dimensionality reduction with remarkable efficiency. In colorectal cancer, Smoother-guided deconvolution revealed plasma cell and fibroblast subtype localizations linked to tumor microenvironment restructuring. Additionally, joint modeling of spatial and single-cell human prostate data with Smoother allowed for spatial mapping of reference populations with significantly reduced ambiguity.

## Background

From subcellular arrangement to tissue compartmentalization, the spatial structure in an organism is highly organized at all scales. This harmonious architecture regulates a diverse variety of biological processes, including embryonic development, neuronal plasticity, and the tumor microenvironment. Recent advances in spatially resolved omics technologies provide unique opportunities to study localization patterns of gene and epigenetic activities, as well as the dynamics of biological systems at cellular and tissue levels(1–5). While existing non-spatial omics analysis methods can be applied to spatial data, the neglect of positional information makes them inadequate in overcoming structured technical noise(6), let alone inferring biologically meaningful spatial organization. Meanwhile, *ad-hoc* spatially aware models often hardcode neighborhood structures into task-specific algorithms(7–15), hindering adaptation to new applications without substantial modification. Even within the same application, models pretrained on one sample usually cannot be applied to another, as the neighborhood graph varies from sample to sample. Additionally, these models are also incompatible with non-spatial data, including the rich single-cell omics atlas datasets, and therefore cannot transfer knowledge learned from non-spatial modalities to enhance spatial analysis.

Physically adjacent spots or cells generally exhibit more similarity compared to distant pairs, with the similarity decaying with distance at different rates (**Supplementary Fig. 1**). Such patterns are consistently observed across tissues, technologies, and modalities — even in single-cell resolution data(**16**) (**Supplementary Fig. 1e**) and in super-resolved tumor microenvironment sections(17, 18) (**Supplementary Fig. 1d and f**). Permutation experiments further confirmed that the long-range similarity structure is not an artifact of contamination or signal bleeding (**Supplementary Fig. 1i**). Despite its universality, spatial dependency in omics data has yet to be formally described in a generic framework independent of downstream applications. Instead, existing algorithms developed for individual tasks often regard spatial variation as a task-specific property, unnecessarily restricting their generalizability to new applications. For example, Markov random field-based models like Giotto(7), BayesSpace(8), CARD(9) and BayesTME(10), although utilizing the same Bayesian message passing mechanism, each introduce structural dependency with unique, integrative, and non-sharable implementations. This practice becomes especially troublesome when the prior belief needs modification, for instance, to encode boundary information or scale to larger datasets. Similarly, graph-based neural networks such as SpaGCN(11) and STAGATE(12) also incorporate spatial structure as a hard constraint and integral part of the model. While interactions between neighbors can be learned adaptively from the data, these models are essentially black boxes, leaving users with minimal control over over-smoothing and signal dilution.

Here we present Smoother, a unified and modular framework for integrating spatial dependency across applications. By representing data as boundary-aware weighted graphs and Markov random fields(19, 20), Smoother explicitly characterizes the dependency structure, allowing information exchange between neighboring locations and facilitating robust and scalable inference of cellular and cell-type activities. Through the transformation between spatial prior and regularization loss, Smoother is highly modularized and ultra-efficient, enabling the seamless conversion of existing non-spatial single-cell-based models into spatially aware versions. We demonstrate the versatility of Smoother by implementing and testing its performance on tasks including cell-type deconvolution and dimensionality reduction. Using simulated and real omics data of different modalities, our benchmarks highlight the substantial advantages of explicitly modeling and incorporating spatial dependencies. Furthermore, Smoother’s soft regularization approach also supports spatially aware joint embeddings of data with and without neighborhood structure, potentially bridging the gap between spatial and single-cell analyses.

## Results

### Overview of the Smoother framework

Smoother differs from existing models in that it treats data-specific dependencies as shared and reusable priors across downstream tasks, encouraging local smoothness on any spatial variable of interest (**Fig. 1**, Methods and Supplementary Notes). Inspired by penalized likelihood methods(21), we decouple the prior belief on dependency from the likelihood of a non-spatial data-generating model. This flexibility allows the same prior to be used in different models, and the same model to accommodate data with varying or even zero spatial structures. In addition, Smoother encodes boundary information that is often neglected in existing Bayesian methods (**Fig. 2**). Specifically, it first builds a spatial graph in which edges connecting physically adjacent locations are scaled and pruned using histological and transcriptomic similarities to remove undesired interactions (**Fig. 2a**). Through graph weighting, users may incorporate additional knowledge from other modalities, even though the actual region boundaries in omics data probably being less distinct (**Fig. 2e** and Supplementary Fig. 2), The spatial graph is then converted into a multivariate normal (MVN) prior with varying degrees of dependencies, along with an equivalent spatial loss that can be appended to non-spatial models (Supplementary Fig. 3. See Methods for detailed recommendations on constructing the prior).

**Figure 1:**
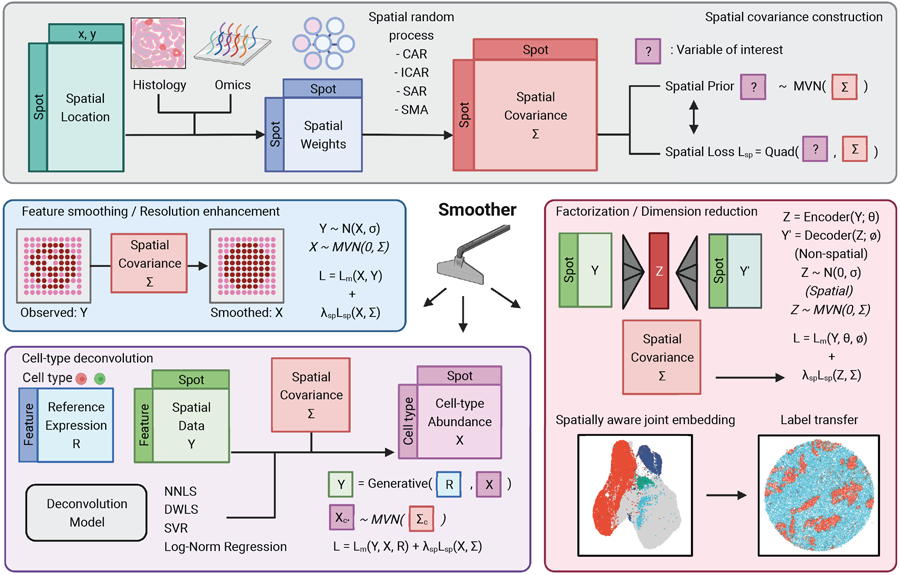
Overview of the Smoother framework. Smoother is a versatile and modular framework designed to incorporate spatial dependencies into various omics data analysis applications. The process initiates with the construction of a weighted spatial graph, derived from physical positions, histology, and additional features, which serves to represent spatial dependencies a priori (top). The prior is subsequently employed as a sparse loss function to regularize spatial variables, such as gene activities, cell-type compositions, and latent embeddings (bottom). Owing to its modular design, the spatial loss can be appended to pre-existing models that were initially developed for non-spatial data, potentially bridging the gap between single-cell and spatial data analysis. The Smoother toolbox includes a selection of spatially aware versions of non-spatial models, including NNLS, DWLS and SVR for cell-type deconvolution, and PCA, SCVI, and SCANVI for dimensionality reduction.

**Figure 2:**
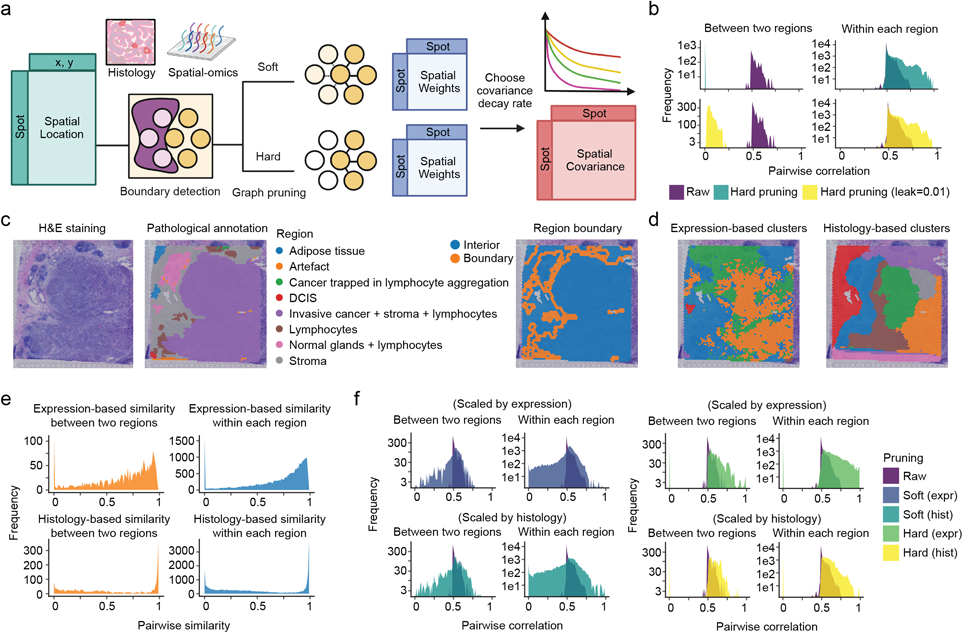
Construction of boundary-preserving spatial priors. **(a)** Schematic illustration of the process for constructing spatial weights and covariance matrices. Smoother first builds a distance-based spatial neighborhood graph, which is subsequently pruned using histology or other features to encode boundary information and inhibit undesired cross-region interactions. **(b)** The impact of hard pruning on the covariance structure. Given the regional membership annotation, hard pruning sets the weights between adjacent spots of different regions to zero (teal, top) or a nominal value (leak=0.01, yellow, bottom), generating distinctive correlation distributions for boundary and interior pairs of spots. **(c)** An example section (1160920F) from the human breast cancer datasets(64). Left to right: H&E staining, manual pathological annotation, and depiction of boundary and interior spots as defined by the pathological regions. **(d)** Unsupervised clustering results of the same slide in **(c).** Spots were clustered based on gene expression (left), or image-based features extracted from the H&E staining image (right). **(e)** The distribution of pairwise feature similarities between boundary pairs from two regions (orange) and interior pairs from the same region (blue), as defined in **(c)** by the pathological annotation. **(f)** The distribution of pairwise correlations imposed by the resulting spatial priors after graph pruning. Left: Soft scaling using expression-(top) and histology-based similarities (bottom). Right: Hard pruning using cluster memberships from expression (top) and histology (bottom) as defined in **(d)**.

As a simple showcase of Smoother, we considered feature smoothing and resolution enhancing, where variables at unobserved locations are inferred from a hidden Markov random field with the MVN prior(22) (**Fig. 1** left and Methods). Unlike other data imputation algorithms(12, 15, 23), our model does not assume feature-level dependencies, making it applicable to a single feature of interest, especially non-expression data. We investigated whether it was possible to overcome data sparsity using spatial context alone. Gene signature scoring is a common approach to evaluate high-level activities of functionally associated gene sets(24), where borrowing information across similar genes may introduce biases and artificially amplify the signal. By penalizing local variations over space, Smoother successfully mitigated dropout effects and improved the separation of localization patterns of cortical layer signatures in a human dorsolateral prefrontal cortex (DLPFC) dataset(25) (Supplementary Fig. 4). Furthermore, Smoother offers ultra-fast functionality, without requiring additional inputs like histological images(15), to enhance spatial resolution to any scale in seconds (Supplementary Fig. 5). While accuracy relies on the assumption that spatially adjacent spots share more similarity, this can be practically helpful when working with shallow sequencing depths. On a Slide-seqV2 dataset of human melanoma brain metastasis(17), we noted improved correlations in the activities of functionally connected modules, such as chemokine and interferon response, after smoothing and enhancing (Supplementary Fig. 6).

### Smoother enhances cell-type deconvolution performance in simulated and real spatial omics data

A common challenge with barcode-based spatial omics technologies that capture a mixture of cells at each location is to disentangle cell-type composition across space. Despite the many deconvolution methods(26–31) developed for spatial transcriptomics (ST) data, few recognize spatial dependency explicitly. Even for these methods, for instance CARD(9) and BayesTME(10), the spatial covariance is hard-coded and not transferable to other models. To fill this gap, we extended four non-spatial deconvolution models using Smoother to resolve the distribution pattern of cell types in a spatially informed manner (**Fig. 1 bottom**). These include non-negative least squares (NNLS), support vector regression (SVR)(32), dampened least squares (DWLS)(33), and log-normal regression (LNR)(34). To benchmark the effect of Smoother, we first simulated ST data with distinct patterns, diverse degrees of spatial heterogeneity, and composition-independent structural noise corresponding to spot bleeding (**Supplementary Fig. 7, 14 and Methods**). In almost every simulation scenario, the inclusion of spatial context consistently yielded more accurate and realistic deconvolution results (**Fig. 3 and Supplementary Figs. 8-14**). Compartment boundaries became notably cleaner as real signals stood out against the prior while noise was smoothed out (**Fig. 3c and g**). This benefit proved highly robust to the selection of models, hyperparameters, and marker genes, while the magnitude shrinks as cell-type-independent spatial variation like spot bleeding grew (**Supplementary Figs. 15-17**). Surprisingly, we observed that CARD failed to take advantage of its own neighborhood modeling, and thus in some scenarios performed worse than a simple non-spatial model. Our results indicate that the parameterization of spatial structure used in CARD is suboptimal, underscoring the importance of a unified and modular framework for dependency representation.

**Figure 3:**
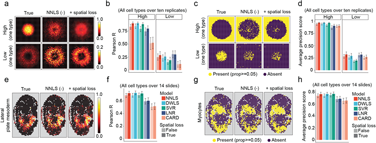
Evaluation of spatial regularization effects on deconvolution accuracy using simulated data. In the first simulation, doughnut-shaped spatial transcriptomic datasets were generated from a scRNA-seq reference(26) on a 50×50 grid. We assigned 15 cell types of high (n=5) and low (n=10) densities to overlapping spatial compartments **(a-d)**. In the second simulation, spatial transcriptomic datasets of the mouse embryo were generated from the sci-Space dataset(63) by pooling single cells of eight types barcoded at the same spot (approximately 200µm pitch) **(e-h)**. **(a)** Relative abundance of two typical cell types in a single replicate. **(b)** Deconvolution accuracy of different methods as measured by Pearson correlation, with results aggregated over ten independent replicates. Error bars denote standard errors of the mean (50 high-density and 100 low-density cell types in total). **(c)** Binary presence status (proportion >= 0.05) of two typical cell types in a single replicate. **(d)** Deconvolution accuracy of different methods, similar to **(b)**, but measured by binary prediction average precision score. **(e)** Relative abundance of the lateral plate mesoderm in one sci-Space slide. **(f)** Deconvolution accuracy of different methods as measured by Pearson correlation, with results aggregated over 14 biological slides. Error bars denote standard errors of the mean (112 cell types in total). **(g)** Binary presence status (proportion >= 0.05) of the myocyte in one sci-Space slide. **(d)** Deconvolution accuracy of different methods, similar to **(f)**, but measured by binary prediction average precision score. Smoother-guided deconvolution models: NNLS (nonnegative least square), DWLS (dampened weighted least squares, modified implementation), SVR (support vector regression), and LNR (log normal regression. The CARD model features its own implementation of spatial regularization.

We next applied all spatially aware deconvolution methods to analyze real spatial omics data of normal, developmental and cancer tissues. Descriptions of each dataset and preprocessing details can be found in the Methods section (**Methods**). We first evaluated deconvolution performance in detecting immune infiltration in a 10x Visium invasive ductal carcinoma dataset(8), where CD3 staining provided ground truth for the presence of T cells. Smoother-guided NNLS faithfully reflected overall T-cell distributions and accurately unveiled the invasive lymphocytic pattern near tumor borders, whereas CARD failed to detect any T-cell signal within the tumor (**Fig. 4a and Supplementary Fig. 18**). The advantage of spatial context modeling was further validated by the elevated correlation of CD3 staining with estimated abundance, outcompeting baseline correlations between gene expression and protein staining (**Fig. 4b and c**). Secondly, to evaluate Smoother’s ability to filter noise while maintaining meaningful boundaries, we performed deconvolution on a 10x Visium dataset of mouse brain(26) and estimated the abundances of 52 neural subtypes across brain locations simultaneously (**Fig. 4d**). As revealed by unsupervised clustering, Smoother notably reduced local discontinuities in cell-type composition, while preserving distinct tissue region boundaries (**Fig. 4e**). In particular, Smoother-based models sharply distinguished excitatory neuron subtypes in cortex and hippocampus, attenuating fluctuations within layers and subregions and producing more precise transitions compared to CARD (**Fig. 4f and g**).

**Figure 4:**
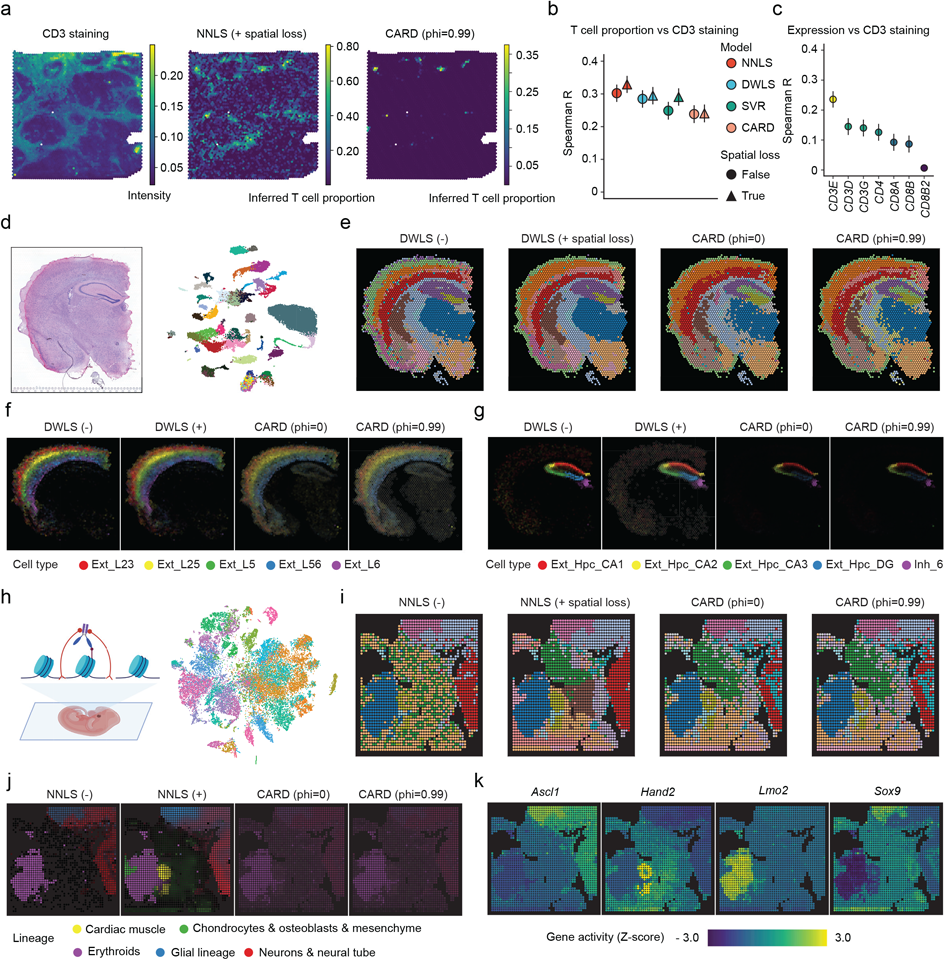
Smoother enhances cell-type deconvolution performance in various spatial omics data. **(a-c)** Detection of tumor T-cell infiltration in a 10x Visium ductal carcinoma sample(8). **(a)** Left to right: Images showing single channel CD3 immunofluorescence staining (FITC/green), T-cell proportions as estimated by Smoother-guided NNLS and CARD, respectively. **(b)** Spearman correlation of predicted T-cell proportions with CD3 fluorescence levels across all spots, with whiskers indicating the 95% confidence intervals. **(c)** Spearman correlation of predicted T-cell proportions with the expression of T cell marker genes across all spots, similar to **(b)**. **(d-g)** Spatial mapping of neurons in the 10x Visium mouse brain section ST8059048(26). **(d)** Left: H&E staining. Right: UMAP visualization of the paired single-nucleus transcriptomic reference of 52 neural subtypes. **(e)** Brain subregions identified by clustering based on estimated cell-type compositions. **(f-g)** Estimated relative abundances of cortical **(f)** and hippocampal **(g)** excitatory neurons. Color intensities are proportional to the estimated cell-type proportion and are scaled by the same factor across methods for visualization purposes. HPC: hippocampus; DG: dentate gyrus (CA4). (**h-k**) Spatial mapping of embryonic cell types in the spatial-CUT&Tag H3K4me3 50µm data of mouse embryo(4). (**h**) Left: Overview of the spatial-CUT&Tag technology, adapted from(4). Right: T-SNE visualization of the scRNA-seq reference of 37 embryonic cell types(35). (**i**) Embryonic subregions identified by clustering based on estimated cell-type compositions. (**j**) Estimated relative abundances of five major lineages aggregated from individual cell types. Color intensities are proportional to the estimated cell-type proportion and are scaled by the same factor across methods for visualization purposes. (**k**) Standardized gene activity of selected marker genes.

Using a spatial-CUT&Tag dataset of mouse E11 embryo, we further assessed the generalizability of Smoother-based deconvolution across modalities (**Fig. 4h and Methods**). Spatial-CUT&Tag(4) profiles chromatin modification using antibodies against histone proteins including H3K27me3, H3K4me3, and H3K27ac. To align the epigenomic data with transcriptomic cell types, we performed deconvolution on epigenomics-based gene activity scores using scRNA-seq reference of 37 embryonic cell types(35) (**Supplementary Fig. 19**). These activity scores differ significantly from expression counts in scale, variability, and biological meaning. Moreover, some transcriptomic cell types may be epigenetically indistinguishable, thereby necessitating more robust deconvolution. Regularization has long been recognized as a solution to the multicollinearity problem(36). Consistently, Smoother salvaged the poor performance of the non-spatial model, producing biologically coherent embryonic compartmentalization (**Fig. 4i**). Close examination of the predicted cell-type composition showed that the Smoother-guided approach excelled in restoring the spatial organization of the embryo with minimal unsolved background (**Fig. 4j**). In particular, only the spatially regularized NNLS model accurately mapped the cardiac muscle lineage to the heart region (**Supplemental Fig. 20-22**). The predicted spatial structures of glial (*Sox9*), neural tube (*Ascl1*), cardiac muscle (*Hand2*), and definitive erythroid (*Lmo2*) lineages were further validated by marker gene activity (**Fig. 4k**).

### Smoother-guided deconvolution unveils distinct localizations of plasma cells and fibroblasts in mismatch repair-proficient colorectal cancer

To further demonstrate the utility of Smoother-guided deconvolution in large-scale cancer ST datasets, we applied the method to a new Stereo-seq mismatch repair-proficient (MMRp) colorectal cancer (CRC) dataset with paired patient-derived scRNA-seq data(18). Utilizing gene modules from prior research(37), we identified eight major cell types and 16 functionally distinct subsets in the scRNA-seq reference, including three plasma cell subpopulations: namely IgG+, IgA+, and IgA+ FOS/JUN+ (**Fig. 5a and b**). *FOS* and *JUN* are B-cell receptor pathway modules and have been shown to be upregulated in tumor-infiltrating B cells in other solid tumors(38).

**Figure 5:**
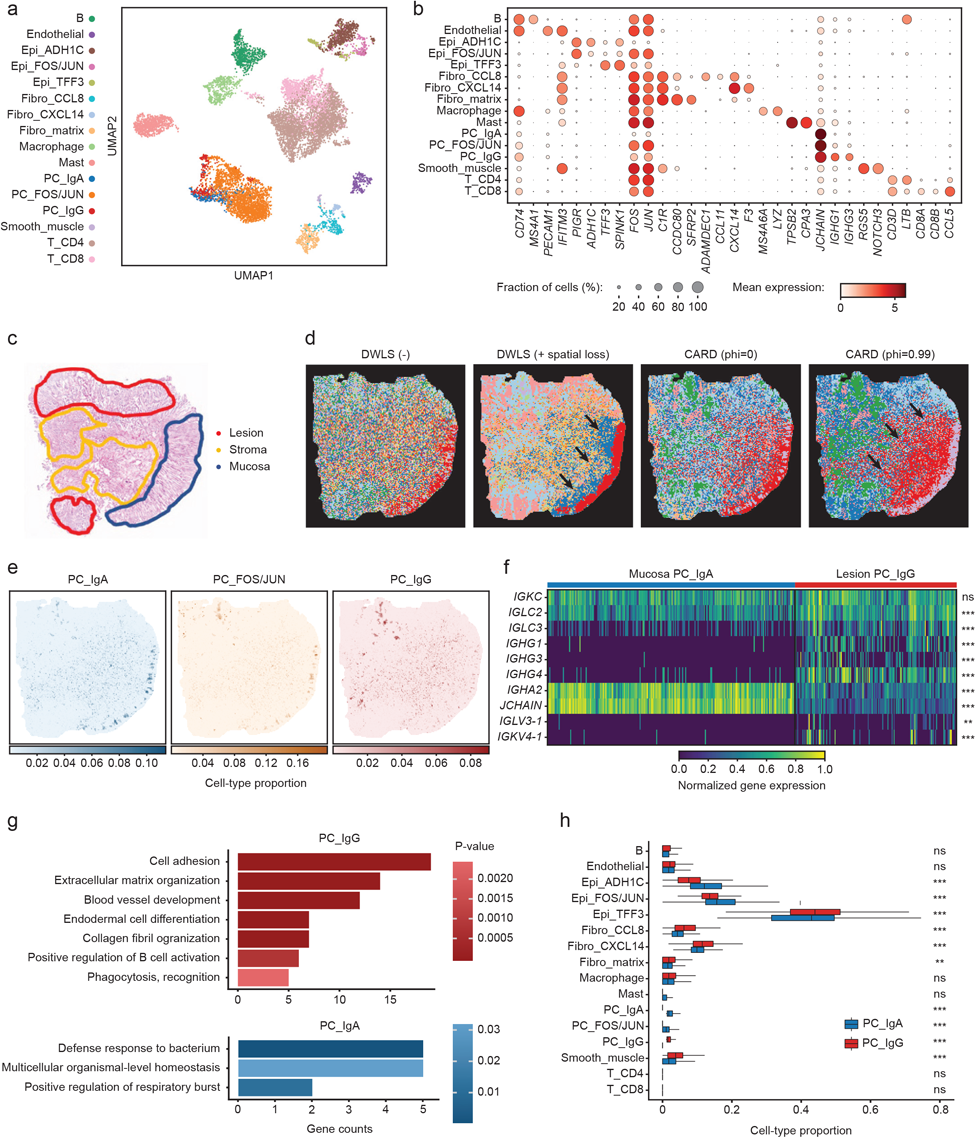
Smoother detects tumor-specific plasma cell subtypes in colorectal adenocarcinoma Stereo-seq slide. **(a)** UMAP representation of the cell types from the patient-derived paired colorectal adenocarcinoma scRNA-seq reference (18). **(b)** Expression dotplot of the top marker genes for each cell type in the scRNA-seq samples. **(c)** Pathology annotation of the histology slide, adapted from(18). **(d)** Slide subregions identified by clustering based on estimated cell-type compositions. Arrows emphasize the mucosa delineation apparent with Smoother but not CARD. **(e)** Spatial visualizations of the cell-type proportions of the three plasma cell subtypes. **(f)** B-cell receptor gene expression heatmap for the IgA plasma cell spots in the mucosa and the IgG plasma cell voxels in the lesion, normalized across spots. **(g)** Colocalization of each cell type with the IgA and IgG plasma cells. **(h)** Gene Ontology pathway enrichment analysis of the IgA plasma cell and IgG plasma cell spots.

We then mapped these single-cell populations to spatial locations through deconvolution. Overall, the Smoother-guided model outperformed CARD in recapitulating the pathological structure, particularly at region boundaries (**Fig. 5c and d**). This facilitated the discovery that the three plasma populations resided in distinct regions: specifically, the IgG+ population were predominantly in the tumor region (lesion) whereas the IgA+ cells were in mucosa (**Fig. 5c and e**). We confirmed the differential localization of IgG+ and IgA+ plasma cells using marker gene expression (**Fig. 5f and Supplementary Fig. 23**). In addition, the lesion IgG+ and mucosa IgA+ spots exhibited divergent B-cell receptor V/C gene usage, with *IGLC2*, *IGLC3*, *IGLV3-1* and *IGKV4-1* all enriched in the tumor, suggesting a difference in the adaptive response between the two plasma cell types (**Fig. 5f**). Gene Ontology pathway analysis further emphasized this difference, with “B-cell activation” enriched in the IgG+ plasma cell sections and “Defense response to bacterium” in the IgA+ plasma cell sections (**Fig. 5g**). Our observation aligns with the established role of IgA+ plasma cells in colorectal mucosal tissues(39), as well as numerous reports of antibody class switching to IgG in the CRC tumor microenvironment(40), which has high potential for diagnostics(41, 42) and therapeutics(37, 43). In stark contrast, CARD incorrectly predicted all plasma cell clones to be IgG+, including in the mucosa region (**Supplementary Fig. 23**).

Additionally, Smoother-guided deconvolution revealed three phenotypically and spatially distinct fibroblast populations, characterized by previously reported ADAMDEC1+/CCL8+, CXCL14+, and matrix transcriptional programs(37), all validated by marker gene expression (**Supplementary Fig. 24**). ADAMDEC1+/CCL8+ and CXCL14+ fibroblasts were preferentially co-localized with the IgG+ plasma cells (**Fig. 5h**), in accordance with the observed presence of CXCL14+ cancer-associated fibroblast and ADAMDEC1+/CCL8+ fibroblast in MMRp CRC(37). Recent work has demonstrated *ADAMDEC1*-driven fibroblastic matrix remodeling in response to inflammation(44), which explains partially the enrichment of collagen and extracellular matrix-related pathways in IgG+ plasma cell sections (**Fig. 5g**). Conversely, CARD again failed to delineate these three fibroblast populations (**Supplementary Fig. 23 and 24**). We repeated the deconvolution analysis on a paired distant normal tissue but observed no differential localizations of plasma cell and fibroblast subpopulations (**Supplementary Fig. 25**).

### Smoother-guided dimensionality reduction and integration of spatial and single-cell data

Learning an informative low-dimensional representation is crucial for understanding the biological dynamics underlying noisy omics data. Smoother’s ability to impose structural dependencies via a versatile loss function allows us to generalize existing non-spatial dimensionality reduction methods to spatial omics data (**Fig. 6a, Methods and Supplementary Notes**). This is further strengthened by the contrastive extension of the spatial loss that better separates distant locations and averts the collapse of embeddings (**Methods**). As a proof of concept, we first developed a spatially regularized principal component analysis (PCA) model(45) and applied it to the human cortex DLPFC data(25). Across all 12 samples, the inclusion of spatial loss consistently improved performance when reconstruction and regularization were balanced, with contrastive loss further strengthening its robustness (**Supplementary Fig. 26a and b**). These enhancements are manifest in the 2D visualization where outer cortical layers are increasingly separated from inner layers and the white matter with stronger regularization (**Supplementary Fig. 26c**). Nevertheless, existing graph-based autoencoders like STAGATE(12) and SpaceFlow(13) achieved even better separation, indicating the intrinsic limitation of PCA as a linear encoder. To dissect the benefit of spatial modeling from simply having a broader parameter space, we replaced the graph attention module in STAGATE with fully connected layers and again incorporated spatial loss during training (**Supplementary Fig. 26d-h**). By increasing the strength of spatial regularization, we recovered 50% - 90% of the performance as measured by embedding consistency (Silhouette score) and clustering accuracy (adjusted rand index) (**Supplementary Fig. 26d-g**). Under 2D UMAP visualization, the space progressively unfolded, and transitions between layers evolved from being non-existent to becoming clear (**Supplementary Fig. 26h**). Together, our results support the potential of spatial loss as a fast and versatile alternative strategy to instill spatial awareness irrespective of model architecture.

**Figure 6:**
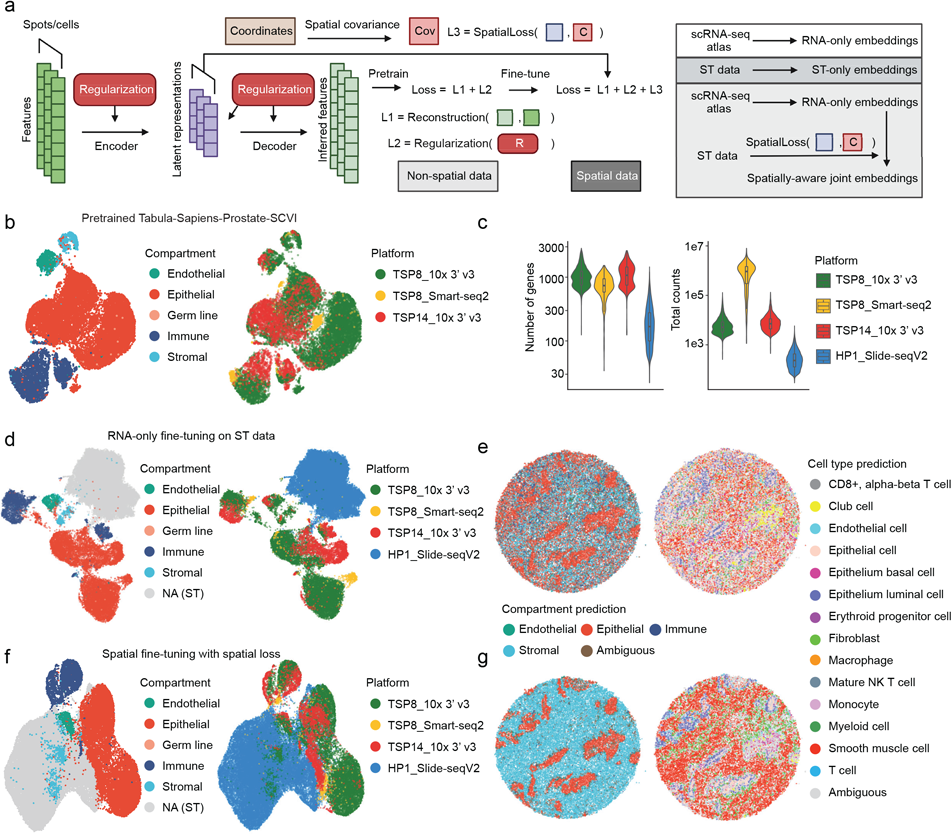
Smoother enables spatially aware joint embeddings of single-cell and Slide-seqV2 data of human prostate and improves reference mapping accuracy. **(a)** Schematic overview of the spatial conversion of non-spatial auto-encoder models in the Smoother framework. Smoother enforces spatial consistency via the detachable loss function. This allows the same model to be trained and applied to both spatial and non-spatial data, generating a joint spatially aware embedding. **(b)** UMAP visualization of the latent representation of scRNA-seq data of human prostate from the Tabula Sapiens(49), colored by tissue compartment (left) and technical batch (right). The representation was generated from a pretrained RNA-only SCVI prostate model(47). **(c)** Violin plots showing the number of expressed gene (left) and total RNA counts (right) per cell or spot in data of different technologies. **(d)** UMAP visualization of the joint latent representation of the Tabula Sapiens prostate scRNA-seq reference and the Slide-seqV2 data of a healthy prostate section(48). Following the SCVI data integration workflow, the RNA-only model was fine-tuned on the query spatial data with unfrozen parameters to mitigate batch effect. **(e)** Spatial visualizations of the tissue compartment (left) and cell type prediction (right) results based on the joint RNA-only embeddings shown in **(d)**. **(f)** UMAP visualization of the joint latent representation generated by SpatialVAE. The spatially aware model has the same architecture as RNA-only models in **(b)** and **(d)**, except it was fine-tuned to minimize the proposed spatial loss in addition to the original reconstruction and KL losses. **(g)** Spatial visualizations of the tissue compartment (left) and cell-type prediction (right) results based on the joint RNA-only embeddings shown in **(f)**.

This modular approach, especially the separation of neighborhood constraints from the autoencoder, offers unique advantages. Through Smoother, we can now adapt pretrained single-cell based non-spatial models to spatial data by fine-tuning over a new spatially aware objective, providing an approach to transfer knowledge from single-cell atlases to spatial omics data (**Fig. 6a**). Specifically, we designed SpatialVAE from SCVI, a prominent deep generative variational autoencoder (VAE) for scRNA-seq data analysis(46, 47). When integrating scRNA-seq datasets, the conventional SCVI workflow includes training a reference model on large atlas datasets, fine-tuning on the query data, and generating joint embeddings of both for label transfer. Here we focused on annotating a human prostate Slide-seqV2 data(48) using a single-cell reference from the Tabular Sapiens(49). After downloading the pretrained model (**Fig. 6b**) from the SCVI model hub, we fine-tuned the model on the spatial data while ignoring coordinates (RNA-only) to remove batch effects. Although SCVI ranks among the top performing data integration tools(50), the Slide-seqV2 data remained distinctly separated from the rest of single-cell data in the joint embedding space (**Fig. 6d**), likely due to its low sequencing depth (**Fig. 6c**). Consequently, most spots were marked as ambiguous, that is, not close to any single-cell clusters within the uncertainty threshold (**Fig. 6e**). In contrast, SpatialVAE’s spatially aware refinements significantly reduced batch effects (**Fig. 6f**), enabling cell label transfer with dramatically reduced ambiguity (**Fig. 6g**). Detailed examination revealed a balanced tradeoff between reconstruction precision and spatial coherence of the embeddings, suggesting that the spatial loss potentially acts via directing the model to focus on spatially consistent features over batch-specific technical noise (**Supplementary Fig.27a**). Using the prostate joint embeddings, we also corrected an annotation mistake in the original publication where cell type labels were mapped to the Slide-seqV2 data using RCTD deconvolution(28). While initially labeled as fibroblasts, the stromal population were indeed primarily composed of smooth muscle cells (referred to as pericytes in(48)) according to SpatialVAE’s prediction (**Supplementary Fig. 27b**). This was confirmed by marker gene expression (**Supplementary Fig.27c-d**).

## Discussion

The inherent neighborhood dependency of spatial omics data motivated us to develop an efficient approach to introduce spatial structure across single-cell applications. Grounded in Bayesian inference and penalized likelihood methods, Smoother imposes regularization at minimal computational burden, particularly by leveraging the sparse nature of the neighborhood graph. In practice, tasks such as enhancing resolution and deconvolving tens of thousands of spots can typically be completed within seconds on a standard personal laptop. The efficiency positions Smoother as an apt solution for the growing demands to explore spatial omics techniques with larger field-of-view, higher resolution, and increased throughput. One caveat to Smoother is that the optimal strength of spatial dependencies, which reflects prior assumptions on the data, is usually agnostic beforehand in downstream applications. Still, we have demonstrated that Smoother is remarkably robust with respect to hyperparameters. It may also be possible to fine-tune the smoothing strength in specific tasks using cross validation and empirical Bayes approaches(51). Collectively, Smoother offers a scalable, and versatile solution to enhance a wide range of tasks including data imputation, deconvolution, and dimensionality reduction. Under this framework, the spatial loss can be melded with any optimization-based non-spatial model, pretrained or otherwise, endowing it with spatially awareness. In light of the ubiquity of structural dependencies, we envision that Smoother may be readily extended to even more applications, such as trajectory inference and cell-cell communication, paving the way for new biomedical discoveries in developmental and disease settings.

## Conclusion

In this study, we introduce Smoother, a powerful and adaptable computational approach designed to integrate spatial structure into omics data analysis. Through spatial priors and losses, Smoother provides a streamlined and efficient way to rewire existing single-cell-based models for spatially informed applications, including feature smoothing, resolution enhancing, cell-type deconvolution, and dimensionality reduction. Benefiting from its robustness, Smoother-regularized deconvolution accurately mapped transcriptomic cell types to spatial epigenomics data. When applied to colorectal cancer sections, it further revealed tumor-associated localizations of plasma cell and fibroblast subpopulations. Furthermore, Smoother’s compatibility with non-spatial data allowed for the spatially informed integration of human prostate data, facilitating cell type prediction with markedly lower ambiguity.

## Methods

### Overview of Smoother

Smoother is a two-step framework that extracts prior dependency structures from positional information and integrates them into non-spatial models for downstream tasks to encourage local smoothness. We use Y_*GXS*_ to denote the spatial omics data of G genes (or the corresponding features in other modalities) at S locations, and L_*FXS*_ the spatial metadata of F spatial features (e.g., image feature extracted from histology) at S locations. The first step of Smoother is to construct a spatial graph G = (V, E, W) where the node set V represents locations and both the edge set E and edge weights W are functions of L_*FXS*_. Then, Smoother computes a covariance structure Σ_*SXS*_ from the graph G and imposes it through the spatial prior on Y_*GXS*_ or other variables of interest. For a comprehensive explanation of Smoother, see below and the **Supplementary Notes.**

#### Construction of spatial priors and losses

Smoother draws from the concept of spatial stochastic process and an extensive body of spatial priors in image processing and geographic data analysis(22, 52, 53). Intuitively, a spatial stochastic process lets us model global dependency characteristics by assuming stationarity and specifying the local interactions of spatial random variables. Here we represent spatial connectivity using a weighted undirected graph G = (V, E, W) where physically adjacent locations V are connected by edges E with varying strengths W. The adjacency matrix w_*SXS*_, which is also called the spatial weights matrix of the underlying spatial process, maps out the connectivity between spots based on physical distance. Here *w_i,j_* = 1 if spot i and j are mutual k-nearest neighbors, otherwise 0. For hexagonal grids like the 10x Visium chip, k is set to 6. To encode domain boundaries, we further scale *W_SXS_* using histological or transcriptomic pairwise similarities (soft-scaling), or manual domain annotations (hard-scaling):

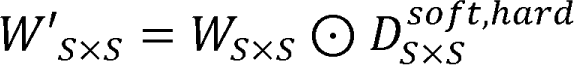

where *D_i,j_^soft^* is the similarity between spot i and j, and *D^hard^_i,j_* is the binary indicator of whether spot i and j belong to the same domain. In practical applications, hard-scaling domains can be defined from histology image segmentation(54), transcriptomic clustering, or expert pathological annotations. For soft scaling, we extract per-spot gene expression or histological features and compute *d_i,j_* as the pairwise similarity in a PCA-reduced space. If scaling by transcriptomics, the first 10 PCs of gene expression and cosine similarity (which is approximately the cosine similarity of Z-scores of full gene expression) are used by default and negative similarities are clamped to zero. If scaling by histology, the feature vector of a spot is the first three PCs of the concatenated RGB values of pixels in the square circumscribed about the spot, and similarity is converted from the Euclidean distance by a Gaussian kernel with band width 0.1 in the normalized PC score space. Specific choices on similarity metrics usually do not have a strong impact on the resulting prior. Empirically, we found the scaling of gene expression to be helpful in maximizing dissimilarity between disparate neighbors.

Subsequently, Smoother translates the spatial weights matrix *W_SXS_* into a covariance structure Σ_SXS_ according to assumptions on the underlying stochastic process (see Supplementary Notes). The covariance is then introduced to any spatial variable of interest, *X_S_*, through a multivariate normal (MVN) prior. For example, in a conditional autoregressive (CAR) process where the graph G describes a Gaussian Markov random field of *X_S_*:

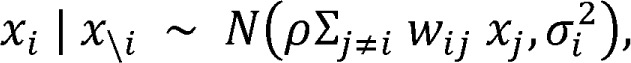

the joint distribution of *X_S_* is a zero-centered MVN distribution with covariance Σ_SXS_, a smoothing prior that can be imposed on *X_S_* in downstream tasks:

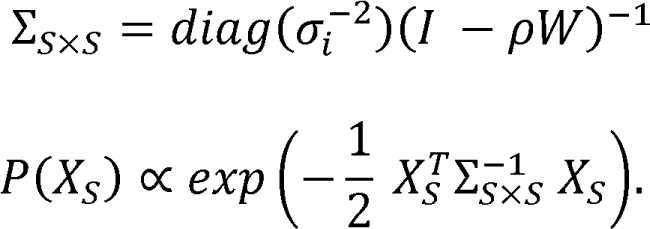

Here ρ is the autocorrelation parameter to make Σ_*SXS*_ positive semi-definite and to control the decay rate of covariance over distance (Supplementary Fig. 3), which can be selected by examining the decay pattern of pairwise similarity (Supplementary Fig. 1). Since Σ_SXS_ is constructed from a boundary-aware graph, the above MVN prior provides a unique channel for neighboring locations to share information while still preserving boundaries.

Smoother offers five different yet related spatial processes: CAR, SAR (simultaneous autoregressive), ICAR, ISAR, and SMA (spatial moving average). Specifically, CAR and SAR are equivalent upon transformation, and ICAR and ISAR are the weights-scaled versions so that the autocorrelation parameter ρ falls in [0, 1]. By adjusting ρ, these models can achieve parallel regularization effects. Based on numerical considerations, we typically recommend using ICAR with varying ρs (or ISAR with smaller ρs) to accommodate data with diverse neighborhood structures. For instance, ‘ICAR (ρ =0.99)’ for data with clear anatomy and ‘ICAR (ρ =0.9)’ for tumor data. SMA is generally not recommended since the resulting inverse covariance matrix tends to be less sparse, potentially slowing down computation.

To render Smoother compatible with existing non-spatial methods, we propose a quadratic regularization loss proportional to the density function of a zero-centered MVN with covariance Σ_*SXS*_:

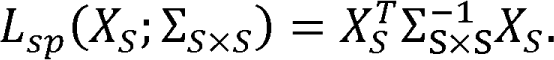

Essentially, for any given model with a loss function L_m_, it is possible to morph the model into a spatially aware version by minimizing a new joint loss function:

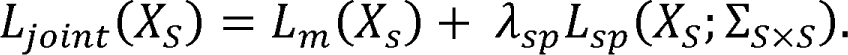

The spatial loss term *L_sp_* regularizes local fluctuations in *X_S_* and makes the inference robust to technical noise. It can be shown that optimizing the new objective is equivalent to finding the maximum a posteriori (MAP) estimator under the spatial prior, and λ_sp_ can be viewed as the strength (inverse variance) of the prior. Most importantly, since *L_sp_* is separated from the model loss, the same model can jointly accommodate data with or without neighborhood structures.

In addition, we implement a contrastive extension of the spatial loss to increase the penalty for pulling distant spots too close, ensuring that the inference does not collapse into trivial solutions. This is done by shuffling spot locations and producing corrupted covariance structures as negative samples,

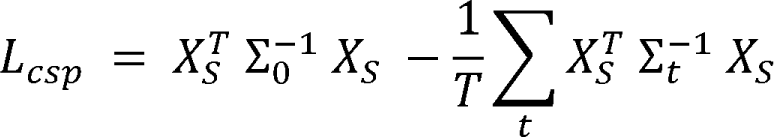

where Σ_0_ is the covariance derived from the correct spatial graph and Σ_*_t_*_, ∈ [1, *T*] are those from corrupted graphs.

#### Data imputation and resolution enhancement

Using local contexts in the prior dependency structure, Smoother is capable of smoothing and imputing spatial omics data at unseen locations for any single spatial random variable of interest *X_S_*. For simplicity, we assume the variable follows a hidden Markov random field model with technical noise being Gaussian iid. This implies that the observation *Y_S_* follows the model:

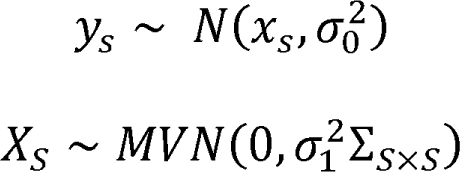

where σ^2^_0_ and σ^2^_1_ are the variance of observation and prior, respectively. Similarly, we can reparametrize the above problem and find the MAP estimator of *X_S_* by solving the following optimization task with a given spatial loss:

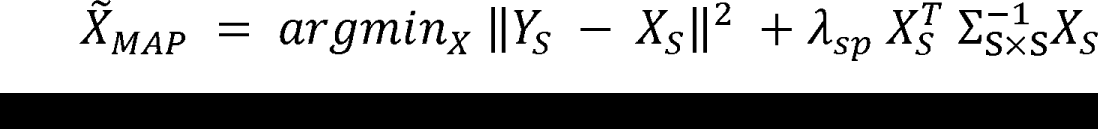

where, λ_*sp*_ < σ^2^_0_/σ^2^_1_ determines the strength of regularization. This is a special case of Tikhonov regularization and is akin to the weighted average filter commonly used for image smoothing. Note that the first L2 term, corresponding to the reconstruction error, can be replaced by other likelihood-based or more sophisticated losses in deep generative models for non-Gaussian variables. When part of the data is missing, the objective function is similar except the first reconstruction term is computed only at observed positions. As Σ_SXS_ is predefined independently of the observation, Smoother can impute the latent value at arbitrary locations and thus increase the spatial resolution.

#### Cell-type deconvolution

We define spatial deconvolution as the problem of inferring cell-type abundances at each location from the observed omics data, with or without cell-type reference information from external data. The task is especially relevant for spatial techniques with limited resolution, including 10x Visium, spatial-CUT&Tag(4) and spatial-ATAC-seq(5), where each profiled location might consists of cells from multiple cell types. The deconvolution model is usually determined by the generative model of the observations *Y_GXS_*. Most deconvolution methods assume a linear relationship between observation and cell-type abundance:

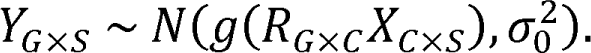

Here, *Y_GXS_* is the observed activities of G features at S spots, *R_GXC_* is the expected reference activities of G features in C cell types, *X_CXS_* is the abundance of C cell types at S spots, σ^2^_0_ is the sampling variability, and g is the data generative function that introduces additional noise such as location-specific biases. Without loss of generality, we follow the same linearity assumption and extend existing deconvolution models to leverage neighborhood information by imposing the spatial prior on the abundance of each cell type:

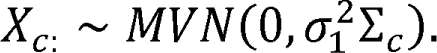

Here, Σ_*C*_ is the prior covariance structure, and 1/σ^2^_1_ represents the strength of prior. When the reference *R_GXC_* is known, usually from a paired single-cell dataset, we solve *X_CXS_* by minimizing the following regularized factorization problem:

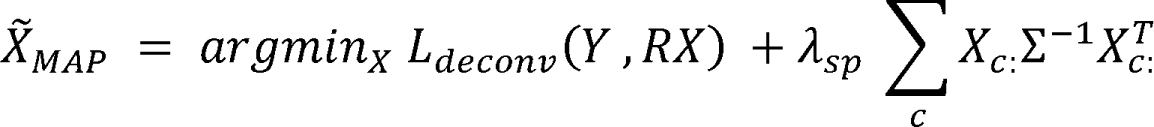

where *L_deconv_*(*Y, Xβ*) is the loss specified by the corresponding deconvolution model. We implemented four spatially aware deconvolution models including non-negative least squares (NNLS), support vector regression (SVR), dampened least squares (DWLS), and log-normal regression (LNR). Further details can be found in the **Supplementary Notes**. When the reference is unknown, the above deconvolution can be solved via matrix factorization, which is also a special case of the dimensionality reduction task, as will be discussed in the next section.

#### Dimensionality reduction

The goal of dimensionality reduction is to infer a condensed low-dimensional representation retaining the data’s essential characteristics. For spatial omics data, latent dimensions should ideally represent continuous dynamics in space. We frame the dimensionality reduction task using a general autoencoder model,

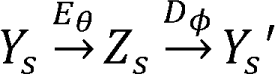

where *E_θ_* is the decoder that projects omics data *Y_S_* ϵ *R^G^* at the location s with G features onto a low-dimensional space *R^H^* with H hidden dimensions, and *D_ϕ_* is the decoder that projects the hidden embedding *Z_S_* ϵ *R^H^* back to the original space.

Using Smoother, we can again impose prior on the hidden embedding to get more coherent representation. For any auto-encoder model with parameter θ, ¢ and reconstruction loss function *L_m_*(*Y_S_*, θ,¢), Smoother regularizes the hidden representation *Z_S_*(*Y_S_*; θ) = *E_θ_* (*Y_S_*) using a spatial loss *L_sp_* and solves a new joint objective

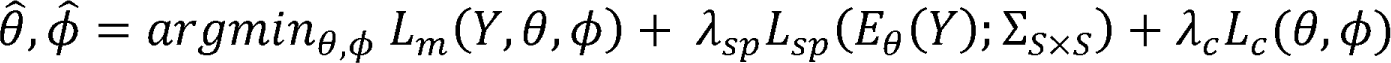

where *L_C_* (θ,¢) is the additional soft constraint loss on model parameters, if any. For linear dimensionality reduction models, including non-negative matrix factorization (NMF)(55), principal component analysis (PCA)(45), and independent component analysis (ICA)(56), the encoder *E_θ_* and decoder *D_<_* are matrix multiplication operators and can be resolved via matrix factorization:

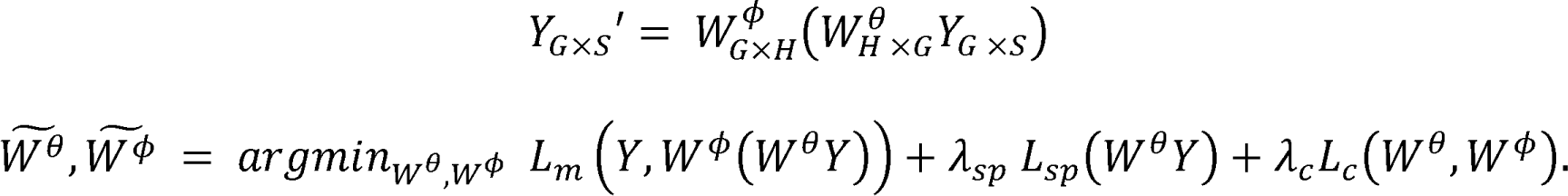

The apparent analogy between the above linear autoencoder model and linear deconvolution suggests that cell-type abundance itself can be viewed as the hidden factor. When the deconvolution reference *R_GXG_* (i.e., *W^ϕ^_GXH_* in the autoencoder) is unknown, factorizing *Y_GXS_* is both a dimensionality reduction task and a reference-free deconvolution task. Further semi-supervised techniques can be applied to maximize the distinguishability of inferred latent states (cell types) using known marker genes.

In this study, we developed corresponding spatial versions of PCA, vanilla deep autoencoder, and Variational Autoencoder (VAE) within the Smoother framework. We assume all hidden dimensions to be independent and regularize them simultaneously. Briefly, the PCA model has symmetric encoder and decoder, requires latent dimensions to be orthogonal and uses L2 norm to measure reconstruction error. The deep autoencoder model removes the symmetry and orthogonality constraints, introduces nonlinearity, and incorporates an orthogonal loss to impose soft constraints on the latent embedding. In a VAE model that describes the data generative process and learns the distribution of latent representations, regularization is by default applied to the mean of the inferred latent distribution.

#### Model implementation

Smoother is publicly available as a Python package (https://github.com/JiayuSuPKU/Smoother/). Models involved in this study are implemented using PyTorch(57) and all optimizations are solved via PyTorch’s gradient-based optimizers (by default Adam for deconvolution and SGD for dimensionality reduction). For convex problems, an alternative Smoother implementation via CVXPY(58) is also available. For VAE models, we adopted the VAE implementation from the Python package scvi-tools(47).

### Preprocess spatial omics data

Unless otherwise noted, we downloaded spatial omics datasets from the SODB database(59) where the data were preprocessed following the Scanpy workflow(60). For data not available through SODB, we used the default Scanpy preprocessing workflow. In **Supplementary Fig. 1**, we used the first 20 PCs of log-normalized expression data to calculate pairwise similarity decay. Preprocessing details of the spatial-CUT&Tag data are provided below in the corresponding subsection. Spatial priors across applications were constructed by default using ICAR with ρ = 0.99 and transcriptomic soft-scaling, unless otherwise specified.

### Recover spatial patterns using Smoother-guided imputation and resolution enhancement

#### Human dorsolateral prefrontal cortex (DLPFC) dataset

For each cortical layer, we calculated gene signature scores using the ‘scanpy.tl.score_genes’ function based on the expression of the top 20 marker genes ranked by log fold change. The spatial prior was constructed under the ICAR model with ρ = 0.99 and λ,_Sp_ = 1. For imputation, we randomly masked out certain proportions of spots in a slide and allowed the target variable to vary in both observed and unobserved locations. For resolution enhancement, we added new spots according to the desired new resolution through midpoint interpolation and ran imputation on the new slide. Variable values at observed locations were fixed during enhancing.

#### Human metastatic melanoma Slide-seqV2 dataset

We preprocessed the Slide-seqV2 data of melanoma brain metastasis (MBM) and extracranial melanoma metastasis (ECM) and the paired scRNA-seq data following the original publication(17) using the R package Seurat(61). Functional signature scores were calculated based on the expression of genes involved in the corresponding pathway using Seurat’s ‘AddModuleScore’ function. We built an ICAR prior with ρ = 0.9 and λ_*sp*_ =1 to impute function scores individually and to enhance the spatial resolution. Pairwise Pearson correlation between functional scores was calculated to evaluate the benefits brought by data imputation.

### Evaluate cell-type deconvolution performance using simulation

#### Simulate ST datasets with spatial patterns

To generate synthetic ST datasets for benchmark, we followed a modified procedure adapted from the cell2location paper(26). These modifications allowed us to: (1) assign arbitrary spatial patterns to zones (co-localized cell-type groups) and (2) introduce additional noise for cell-type-independent dependencies (e.g. spot bleeding). In brief, we initially overlaid designated patterns with a two-dimensional Gaussian process to generate the per-zone abundance values in space, then assigned cell types to these patterns. Next, we sampled gene expression profiles at each location from a scRNA-seq reference according to the simulated cell-type composition. The ST data is further combined with various sources of noise, including lateral diffusion where each spot shares a certain proportion of mRNA to its neighbors directly, and multiplicative per-gene gamma noise (by default shape = 0.4586, scale = 1/0.6992) for sampling error. The simulation code is provided as a stand-alone tool within the Smoother package to facilitate future benchmarking on related tasks.

#### Model comparison and evaluation

In the study, we focused our benchmarking on CARD, the only published spatially aware deconvolution method shown to be superior to other existing methods(9). However, Smoother is compatible with different deconvolution strategies. Any method, including the DWLS(33), nu-SVR(32), and LNR(34) re-implemented in this study, may be seamlessly transformed to take advantage of neighborhood information. CARD is a non-negative linear factorization model that introduces spatial dependencies through a conditional autoregressive prior on cell-type abundances, which is mathematically similar to the Smoother-guided NNLS model. We adhered to the tutorial and ran CARD with default parameters. In certain simulation scenarios, a truncated reference signature matrix with fewer genes was used as input, bypassing CARD’s internal reference construction process. This was to reflect potential discrepancies between the external reference and observed ST data, and to separate the impact of reference quality from algorithm performance. For the non-spatial CARD model, we set ρ to zero, effectively disabling any spatial interactions.

The performance of deconvolution depends on the distribution properties of the input data. For example, RNA-seq counts are typically skewed and, if left uncorrected, can bias the estimation against lowly expressed genes and rare cell types(34). One practical solution is to perform deconvolution on the logarithmic scale, i.e., replacing Y and x with 1ay1p(Y) and 1ay1p(x). Although this approach is not physically sound, it has been shown to significantly improve model performance, so does the square root scaling √Y and √x albeit to a less extent (data not shown). For benchmarking purposes, unless otherwise stated, we supplied log-scale ST data to NNLS, DWLS, and nu-SVR and raw-scale ST data to LNR and CARD. The reference expression matrix was computed by averaging the normalized expression of marker genes across all cells of a given cell type from an external scRNA-seq dataset, followed by log-transformation where necessary. For more general cases, such as in epigenomics deconvolution where the reference and observation data may not be on the same scale or even from the same modality, users may include whatever preprocessing steps that best fit the dataset.

For Smoother-guided models, we constructed the spatial prior using ICAR (ρ = 0.99, which was also the optimal ρ in CARD) and scaled the graph using transcriptomic similarity. The strength of the spatial prior λ,_*sp*_ is set to 0 for all non-spatial baseline models, 1 for spatially aware NNLS, SVR, LNR, and 3 for spatially aware DWLS to adjust for the inflated model loss after scaling. This is not necessarily the best performing setting as revealed in the parameter sensitivity analysis (**Supplementary Fig. 15-17**). Nevertheless, we fixed the strength across all benchmarks since the benefit of spatial context is rather robust. Cell-type abundances were set to be non-negative in all models and were normalized to output the final cell-type proportions at each spot.

#### Benchmark deconvolution performance using the simulated doughnut-shaped data

We obtained the single-nucleus RNA-seq data of mouse brain (5705STDY8058280, 5705STDY805828) along with cell type annotations for each cell from https://www.ebi.ac.uk/arrayexpress/experiments/E-MTAB-11115/, and only kept the 15 most abundant neural subtypes for simulation. The dataset was further split into two, one to simulate ST data and the other as the deconvolution reference. We generated ST datasets on a 50×50 grid containing five high-density (average of four cells per occupied spot) and 10 low-density (0.4 per occupied spot) cell types. Each cell type was assigned to a unique but overlapping doughnut-shaped distribution pattern. Cell-type-independent spatial dependencies were introduced by sharing 0/10%/50% of mRNA counts at each location to its first-degree neighbors (i.e. spot bleeding). For each simulated dataset, we performed deconvolution using four sets of genes: (1) The union of the top 20 marker genes for each cell type (n=283), (2) the union of the top 50 marker genes (n=656), (3) all discriminative informative markers genes whose log2(fold change) is larger than 1 in one and only one cell type (n=2693), or (4) informative genes selected by the CARD model (n=7972). This in total brings 12 scenarios, each is replicated 10 times. All preprocessing and differential expression analysis steps were performed using the Python package Scanpy(60).

We evaluate deconvolution performance using three metrics: (1) The mean square error and (2) the Pearson correlation between ground truth and the estimated proportion per cell type, and (3) the average precision score of binary prediction of whether a cell type is present at a given location (abundance >= 0.05) using the Python package Scikit-learn(62). Results across cell types and experiments (replicates) are aggregated together for boxplot and barplot visualizations.

#### Benchmark deconvolution performance using the sci-Space data of mouse embryo

We obtained the sci-Space data(63) of 14 mouse embryonic slides from GEO under the accession number GSE166692. Each single cell in the sci-Space data is labeled with an approximate spatial coordinate (∼200 µm pitch). Cells of low-quality or from rare cell types (less than 100 cells in any slide) were filtered out. For the remaining eight cell types, we averaged the expression of 298 marker genes defined in the original paper to compute the deconvolution reference. To generate ST data of the same resolution, we pooled together all cells with the same barcode and resampled cells in each spot to adjust for the uneven density of captured cells. The actual cell count at each location is determined by a gamma-Poisson distribution with an expectation of 10. Since most coordinates have only one associated cell, the resampling was executed with replacement from both the current spot and adjacent locations (with half probability), effectively increasing the diversity at each spot. To further challenge the deconvolution, we down-sampled the synthetic ST data to a maximum of 5,000 UMI per spot and added multiplicative per-gene gamma noise with the mean around 0.35.

### Evaluate cell-type deconvolution performance using real spatial omics datasets

#### Benchmark deconvolution performance using the breast tumor dataset with staining

We obtained the invasive ductal carcinoma (IDC) data(8) with DAPI and anti-CD3 staining from 10x Genomics at https://support.10xgenomics.com/spatial-gene-expression/datasets and performed deconvolution to evaluate T-cell infiltration. The reference gene expression of nine cell types, including one tumor epithelial type, was computed using a scRNA-seq atlas of 26 breast cancer samples(64). For Smoother-guided models, we selected the top 20 unique differentially expressed genes for each cell type using Seurat(61) (180 genes in total) as the input for deconvolution. For CARD, we provided the scRNA-seq data with all genes as input and run with default parameters. Performance was first evaluated by measuring the Spearman correlation between the CD3 immunofluorescence (IF) intensity and the T-cell deconvolution proportion over all spots on the slide. We then investigated if these models could reliably reveal T-cell infiltration in tumor regions (labeled as invasive carcinoma regions in the original publication), a question of paramount importance for immunotherapy research. Specifically, we fitted a two-component Gaussian mixture model on the CD3 intensity and predicted 377 out of the 2078 spots in the tumor regions to be T-cell positive. Results were visualized as pie charts in **Supplementary Fig. 18**.

#### Visualize the smoothing effect on region boundaries using the 10x Visium mouse brain data

We obtained the 10x Visium dataset of mouse brain(26) and the paired snRNA-seq reference from ArrayExpress under accession IDs E-MTAB-11114 and E-MTAB-11115. The reference expression matrix for deconvolution was calculated for 52 neural subtypes after removing seven unknown or low-quality cell types. For Smoother-guided methods, the union set of the top 20 marker genes for each cell type discovered by Scanpy(60) was used as the input for deconvolution. For CARD, all QC-passed genes were supplied, and the reference matrix was estimated by CARD. To visualize brain regions with distinct cell-type compositions, we first built a KNN graph based on the the first 20 PCs of inferred cell-type proportions, then applied Leiden clustering using Scanpy(60) and aligned the resulting clusters across methods via linear sum assignment. Clustering resolution was manually adjusted for different deconvolution methods so that the number of clusters generated per method was approximately the same.

#### Spatial mapping of scRNA-seq defined cell types to the spatial-CUT&Tag data of mouse embryo

We obtained the spatial-CUT&Tag data and preprocessing scripts from the original publication(4). Specifically, the R package ArchR(65) and the ‘getGeneScore_ArchR.R’ script were used to compute gene activity scores for the top 500 variable genes using the same set of parameters. We removed spurious spots with outlier average gene activity (more than three standard deviations away from the mean), which lined up in one column or row on the grid and were considered as technical artifacts. The remaining spots were used in all downstream analyses.

For deconvolution, scRNA-seq data of E11 mouse embryo was obtained from Mouse Organogenesis Cell Atlas (MOCA)(35) and processed as described in the spatial-CUT&Tag paper. Specifically, we used the script ‘integrative_data_analysis.R’ to read and preprocess the subsampled (100k cells) data and Seurat’s ‘FindVariableFeatures’ function to identify the top 500 variable genes. We then computed the pseudo-bulk expression of variable genes for cell types annotated in ‘Main_cell_type’ group and used it as the deconvolution reference. Spatial domains of cell types were determined and visualized using the same clustering strategy developed in the mouse brain analysis above.

### Deconvolution analysis of the Stereo-seq data of mismatch repair-proficient colorectal cancer

#### Re-analyze and annotate the paired scRNA-seq data

We downloaded processed Stereo-seq and paired scRNA-seq data(18) from CNGB Nucleotide Sequence Archive under accession ID CNP0002432. The data was further processed as described below using Scanpy. After filtering out outliers (‘n_genes_by_counts >= 2500’ or ‘pct_counts_mt’ >= 5), we first clustered the single-cell reference into eight major cell types (T cell, B cell, mast cell, macrophage, plasma cell, epithelial cell, endothelial cell, and fibroblast) and confirmed their identities via marker genes. To identify functional subtypes, the three most abundant populations (T cells, plasma cells, and epithelial cells) as well as fibroblasts were separately re-clustered after regressing out the effect of sample donor. Each subtype was annotated according to the gene modules reported in (37). In particular, we re-discovered three plasma cell subsets (IgG+, IgA+, and IgA+ FOS/JUN+), three epithelial subsets (ADH1C, FOS/JUN, and TFF3), and three fibroblast subsets (CCL8, CXCL14, and Matrix), all with clear markers. These subpopulations were either missed or misclassified in the original publication (18), likely due to batch effects.

#### Deconvolute Stereo-seq tumor and normal sections of patient P19

We extracted raw counts of the bin100 (50µm x 50µm) Stereo-seq data from the tumor and the paired distant normal tissue samples of patient P19 for separate deconvolution. For Smoother-guided deconvolution, the reference profile was the log averaged expression of the top 50 marker genes for the 16 subtypes (695 genes in total). For CARD, the reference was based on the same set of genes without log-scaling the expression. The default CARD-constructed reference contained much more genes (15025 genes in total) and yielded worse results (data not shown). The ‘DWLS (+ spatial loss)’ model shown in **Figure 5** was regularized using the ICAR prior with ρ = 0.99 and λ,_*sp*_ = 3. Spatial domains of cell-type proportions were determined and visualized using the same clustering strategy (i.e. leiden clustering after PCA transformation) described in the mouse brain analysis above.

#### Differential localization of the IgG+ and IgA+ population in the tumor section

We divided spots into IgA- and IgG-specific groups based on the log ratio between ‘PC_IgA’ and ‘PC_IgG’ proportions estimated from the ‘DWLS (+ spatial loss)’ model (log ratio thresholds: >0.5 and <-0.5). For robustness, noisy proportions less than 0.01 were removed and a pseudo-count of 0.01 was added to all spots when calculating the log ratio. We then computed differentially expressed genes between the two groups (Wilcoxon test, p<=0.01) and further identified enriched ‘GO:BP’ pathways in each region using the function ‘scanpy.queries.enrich’. For colocalization analysis, we again removed noisy proportions (<0.01) and calculated statistical significance using the Wilcoxon test.

### Joint embedding of Slide-seqV2 human prostate data and the Tabula Sapiens prostate scRNA-seq reference using SpatialVAE

We acquired raw Slide-seqV2 count data(48) of healthy human prostate samples and the associated annotations from https://github.com/shenglinmei/ProstateCancerAnalysis. The pretrained reference SCVI model and the training scRNA-seq data from the Tabula Sapiens prostate were downloaded from SCVI model hub at https://huggingface.co/scvi-tools/tabula-sapiens-prostate-scvi. Adhering to the conventional SCVI data integration workflow, we first fine-tuned the pretrained model on the Slide-seqV2 data with ‘unfrozen=True’ for 100 epochs to mitigate batch effects. This step also updated the latent representation of the single-cell reference. Model convergence was confirmed by inspecting the evidence lower bound (ELBO). For spatially informed fine-tuning, we converted the SCVI model into a new SpatialVAE model with spatial loss (ICAR, ρ = 0.99, λ,_*sp*_ = 0.01). The strength λ,_*sp*_ was selected to balance the spatial loss and the reconstruction accuracy (measured by log likelihood). A SpatialVAE model shares the same architecture and initial model parameters with the baseline model, which can be either the Tabula Sapiens reference model or the RNA-only fine-tuned model. In the study, we initialized SpatialVAE from the updated model and further fine-tuned with respect to the new objective for another 100 epochs. This was mainly to highlight the tradeoff between reconstruction accuracy and spatial consistency. Skipping the RNA-only fine-tuning step will not affect the performance of the final spatial model. To transfer cell labels, we followed the SCVI reference mapping workflow described in (66). Briefly, for each query spot we first identified nearest neighbors in the reference, then assigned labels to the spot based on annotations of the neighbors. The prediction uncertainty score was calculated based on the pairwise distance between the query and its reference neighbors in the latent space. Spots with uncertainty >0.2 were labeled as ‘ambiguous’.

### Evaluate dimensionality reduction performance using the DLPFC dataset

All models employed expression of the top 2000 highly variable genes in each of the 12 DLPFC samples(25) as input for dimensionality reduction. For PCA, the expression was further scaled after log-normalization. Based on the ground truth layer annotation, we evaluated the quality of the obtained latent representation using two metrics: embedding consistency measured by the Silhouette score using Scikit-learn(62), and clustering accuracy measured by adjusted rank index where we clustered spots into the observed actual number of regions (cortical layers) using the R package ‘Mclust’(67). The STAGATE model was configured with two graph attention layers of 128 and 30 units (i.e., 30 latent dimensions) and trained using default parameters. The SpaceFlow model had two fixed graph convolutional layers. We set the hidden dimension size ‘z_dim’ to 30 and again trained the model using the default setting. The baseline vanilla deep autoencoder contained two fully connected layers of dimension 128 and 30, with batch normalization and ‘ELU’ as the activation function. For Smoother-guided models, we constructed the regular spatial loss using the ICAR model and ρ = 0.99. To construct the contrastive loss, we generated 20 corrupted graphs and set the relative importance of negative samples to 0.05. When training the deep autoencoder, we also introduced an additional orthogonal loss regularizing the latent space to prevent embedding collapse.

## Declarations

### Ethics approval and consent to participate

Not applicable.

### Consent for publication

Not applicable.

### Availability of data and materials

The python package Smoother developed in this study, tutorials, and the scripts used to simulate ST data are publicly available at https://github.com/JiayuSuPKU/Smoother/ under the BSD 3-Clause License. All additional analysis codes for reproducing results and figures presented in this study will be made available as a separate GitHub repository upon the publication of this work. Descriptions and links to all public datasets analyzed in this study can be found in the corresponding Methods sections. These include the single-nucleus RNA-seq data and the 10x Visium data of mouse brain(26) downloaded from ArrayExpress (https://www.ebi.ac.uk/arrayexpress/experiments/) under accession IDs E-MTAB-11114 and E-MTAB-11115, the sci-Space mouse embryonic data(63) downloaded from GEO at https://www.ncbi.nlm.nih.gov/geo/query/acc.cgi?acc=GSE166692, the 10x Visium invasive ductal carcinoma (IDC) data(8) with DAPI and anti-CD3 staining from 10x Genomics at https://support.10xgenomics.com/spatial-gene-expression/datasets, and the spatial-CUT&Tag mouse embryonic data(4) downloaded from GEO at https://www.ncbi.xyz/geo/query/acc.cgi?acc=GSE165217 with preprocessing scripts from Zenodo at https://zenodo.org/record/5797109#.Y9wezOyZP0p. The CRC Stereo-seq and paired scRNA-seq data(18) were obtained from CNGB Nucleotide Sequence Archive under accession ID CNP0002432 at https://db.cngb.org/search/project/CNP0002432/. The human prostate Slide-seqV2 data (48) were obtained from https://github.com/shenglinmei/ProstateCancerAnalysis.

## Authors’ contributions

J.S. conceived the original idea and developed the computational framework. R.R. and D.A.K. supervised the project. J.S., J.R., X.F., G.Z., and R.S.E carried out the analysis. J.J developed the data simulation scripts. Y.W. and B.I. contributed to the analysis of MBM data and the interpretation of the results. L.A. contributed to the analysis and interpretation on human prostate cell-type prediction. J.S. took the lead in writing and revising the manuscript with input and critical feedback from all authors. All authors read and approved the final manuscript.

## Acknowledgements

The authors thank Y. Choi for contributions in benchmarking, P. Sims, Y. Shen, and C. Zhang for early discussions, and members of the Rabadan lab and Knowles lab for helpful comments and feedback throughout the project.

