## Supplementary Figure for "Smoother: A Unified and Modular Framework for Incorporating Structural Dependency in Spatial Omics Data"

### Supplementary Figure 1

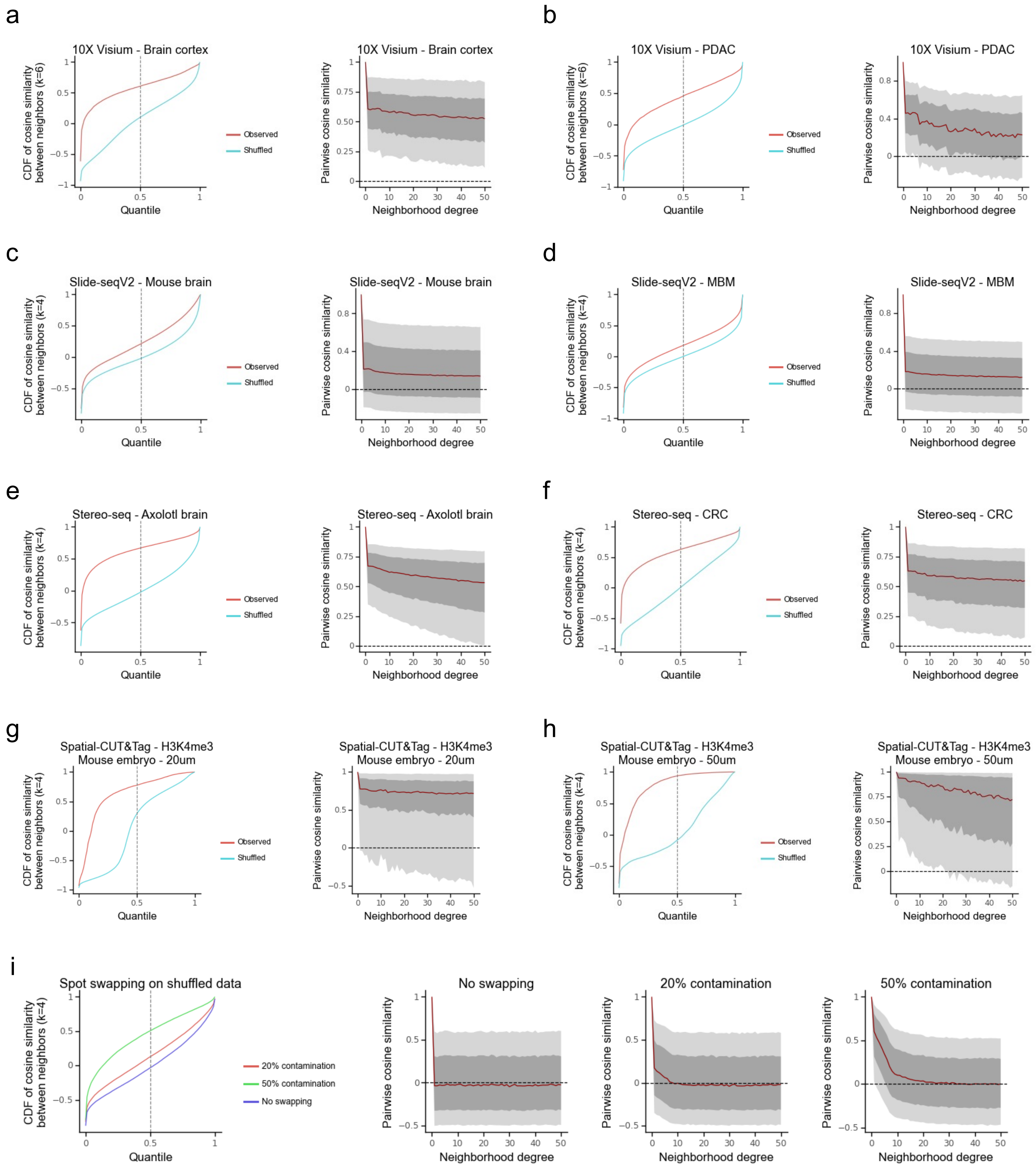


**Supplementary Figure 1: Spatially adjacent spots share similar profiles, a common property of spatial omics data observed across biological systems and technologies.**

The figure demonstrates the distributions of pairwise cosine similarity between neighboring spots in spatial omics data. The similarity is calculated based on the first 20 PCs of log-normalized gene counts (for transcriptomics data) or gene activity scores (for epigenomics data). All transcriptomics data were preprocessed following the standard Scanpy workflow(1) by SODB(2). **Left in each panel:** Cumulative distribution of pairwise similarity between k-nearest physically adjacent spots. Red line shows the observed distribution whereas blue line shows the null distribution of random shuffled spots. **Right in each panel:** Variability in pairwise similarity as a function of neighbor distance. For each neighborhood degree level k (x-axis), pairwise similarities were calculated between the k-th nearest neighbors (self-similarity of 1 when k=0). The light gray area indicates the 10%-90% range, dark gray the 25%-75% range, and the red line the median, respectively. **Datasets inspected:** (a) 10x Visium human brain dorsolateral prefrontal cortex (DLPFC)(3). (b) 10x Visium human pancreatic ductal adenocarcinoma (PDAC)(4). (c) Slide-seqV2 mouse hippocampus(5). (d) Slide-seqV2 human melanoma brain metastasis (MBM)(6). (e) Stereo-seq axolotl brain (segmented and binned at single-cell resolution)(7). (f) Stereo-seq human colorectal cancer (CRC, binned into 50 μm x 50 μm spots)(8). (g) Spatial-CUT&Tag mouse embryo, H3K4me3 20μm(9). (h) Spatial-CUT&Tag mouse embryo, H3K4me3 50μm(9). (i) Shuffled Stereo-seq single-cell resolution data. Per contamination rate p, each spot contains 1-p of its original RNA counts and p of RNA counts from its adjacent neighbors.

Supplementary Figure 2

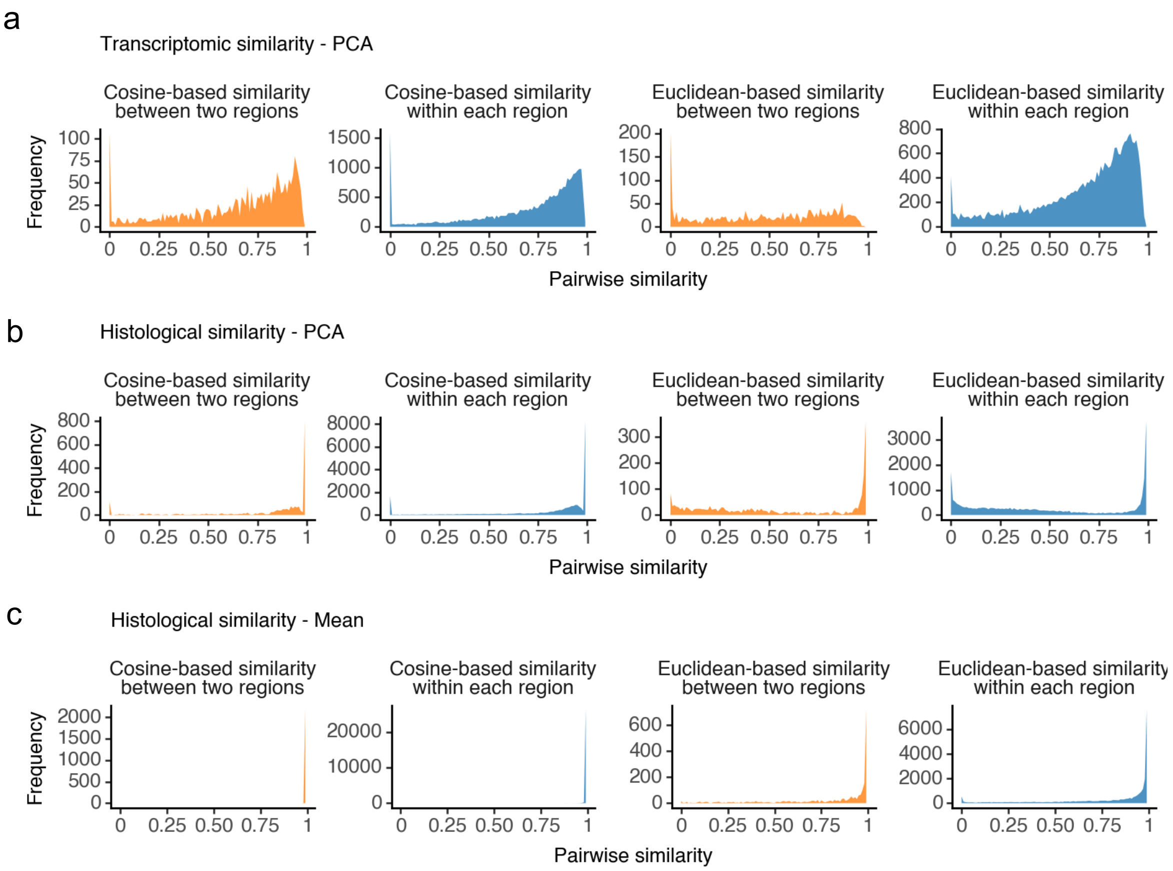


**Supplementary Figure 2: Distributions of pairwise similarities between boundary pairs from two regions and interior pairs from the same region, related to Figure 2e.**

Boundary pairs from two regions (orange) and interior pairs from the same region (blue) are defined in **Figure 2c** by the pathological annotation. (a) Transcriptomic similarities are calculated based on the log-normalized expression data of the top 2000 highly variable genes. PCA with zero-centering and scaling was applied to reduce the feature to the first 10 PCs. Left two panels: Cosine similarity. Right two panels: Euclidean similarity as converted from the Euclidean distance using an exponential kernel of bandwidth 0.01. (b) Similar to (a), except the feature is the concatenated RGB values of pixels of the H&E staining covered by the spot. PCA with zero-centering and scaling was applied to reduce the feature to the first 10 PCs. (c) Similar to (b), except the feature is the RGB values averaged over pixels of the H&E staining covered by the spot.

Supplementary Figure 3

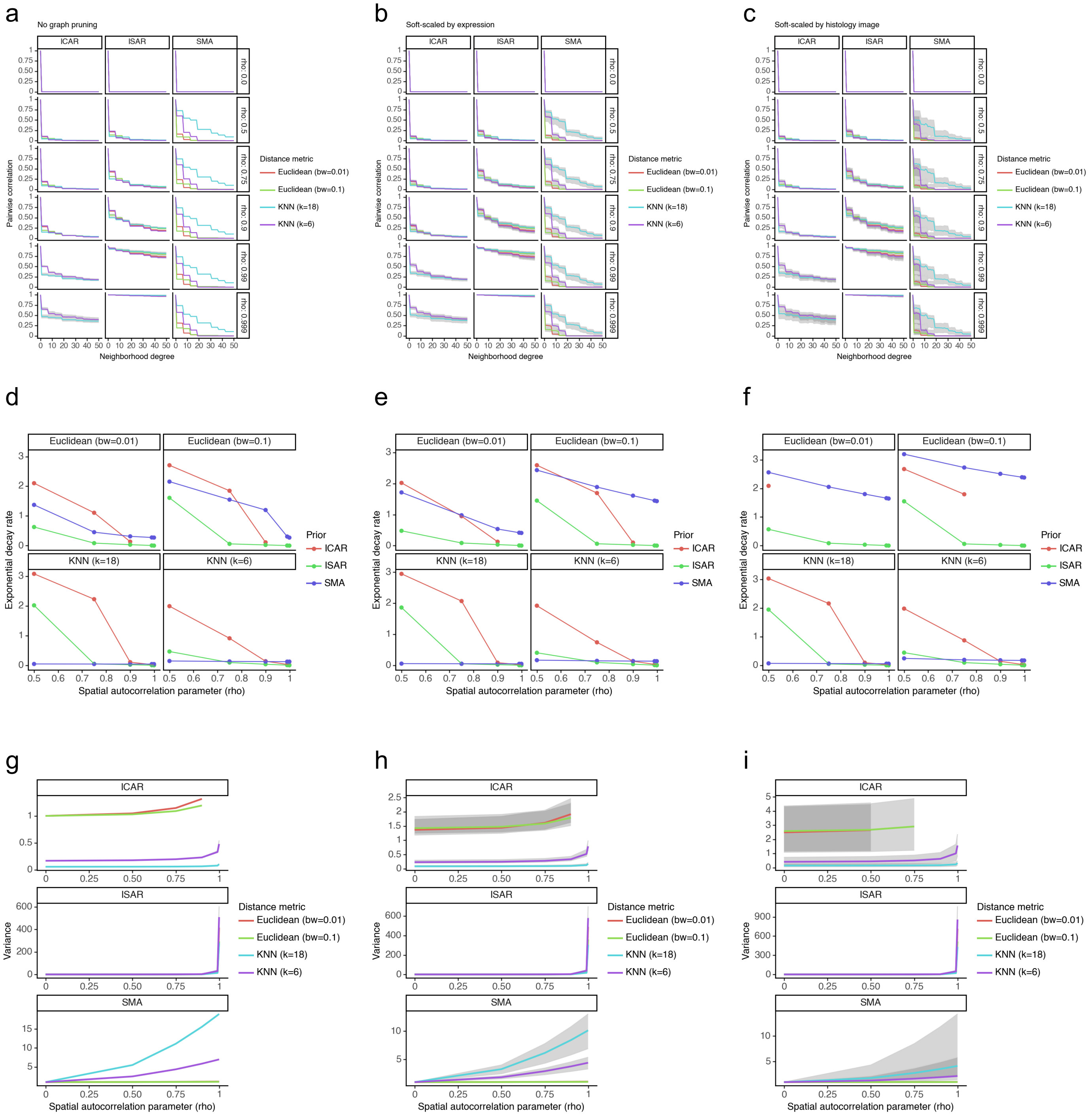


**Supplementary Figure 3: Covariance structure analysis under different spatial random processes and autocorrelation parameters.**

(a-c) Distribution of pairwise correlation between the k-th nearest neighbors, as specified by the spatial stochastic process (columns) and the autocorrelation parameter (rows) ‘rho’. Intuitively, the autocorrelation parameter ‘rho’ can be viewed as the proportion of information that comes from the neighbors. The distributions are shown with colored lines and the associated gray areas indicating the median and 25%-75% range of correlation, respectively, over all neighboring pairs of the same neighborhood degree k (self-correlation of 1 when k=0). Color denotes different strategies to construct the spatial weights matrix. The spatial weights in the neighborhood graph are further scaled by expression similarity in (b), and by histology similarity in (c). Estimated exponential decay rates of median correlation are depicted in (d-f), and distributions of per-spot variances are shown in (g-i).

### Supplementary Figure 4

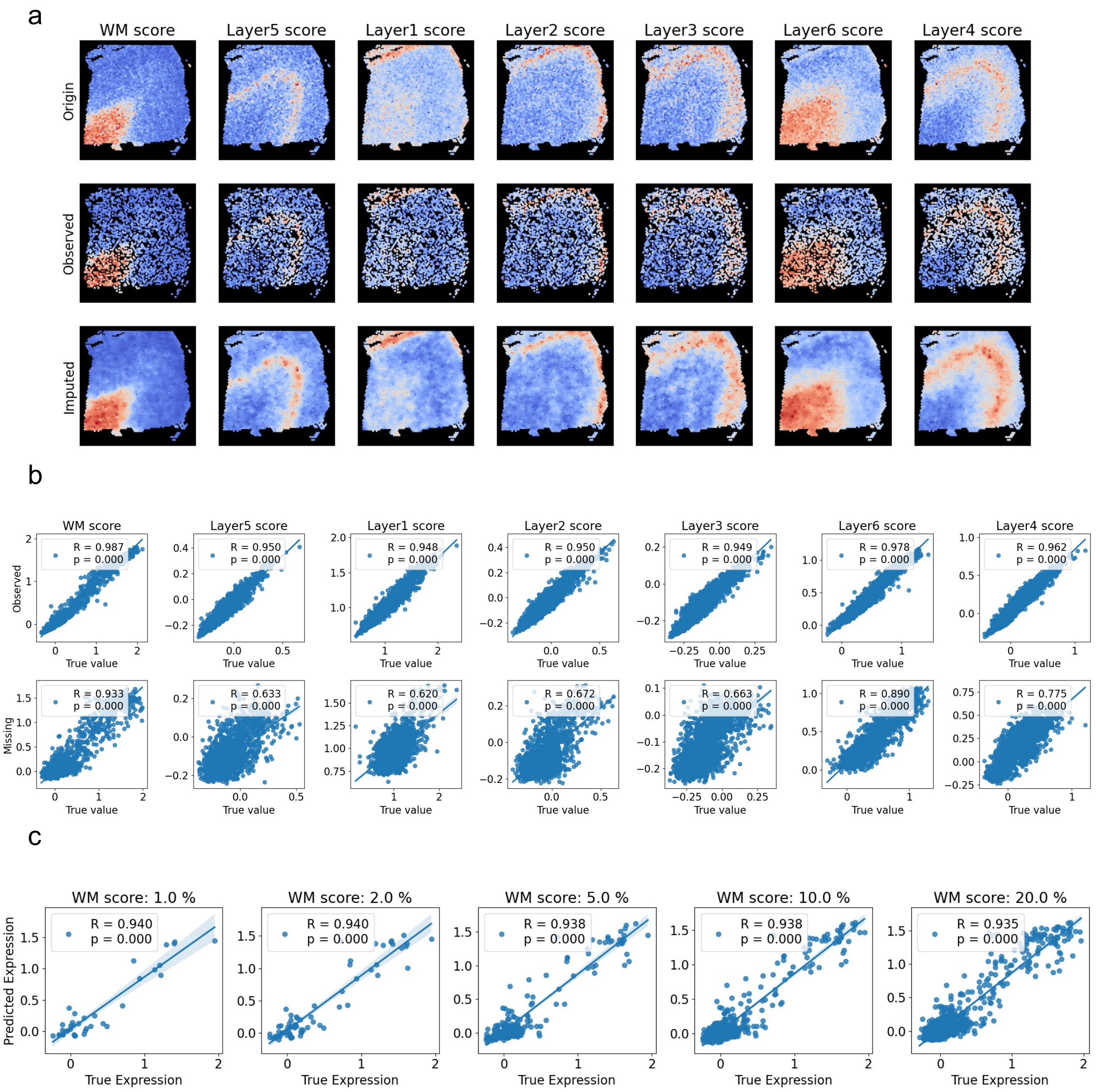


**Supplementary Figure 4: Application of Smoother for gene signature score imputation and smoothing in the DLPFC dataset (151673).**

Gene signature scores were defined per region using 'scanpy.tl.score_genes' as the weighted scaled average of top20 marker genes of that region (adjusted for gene background). (a) Recovery of spatial patterns of signature scores through Smoother-guided data imputation. 2000 out of total 3639 spots are supplied as inputs (second row) to infer the score in all spots (third row). (b) Scatter plots of the imputed score against the observed score in input spots (top) and masked-out spots (bottom). (c) Imputation performances of the white matter (WM) score at masked-out locations as a function of mask-out rate.

### Supplementary Figure 5

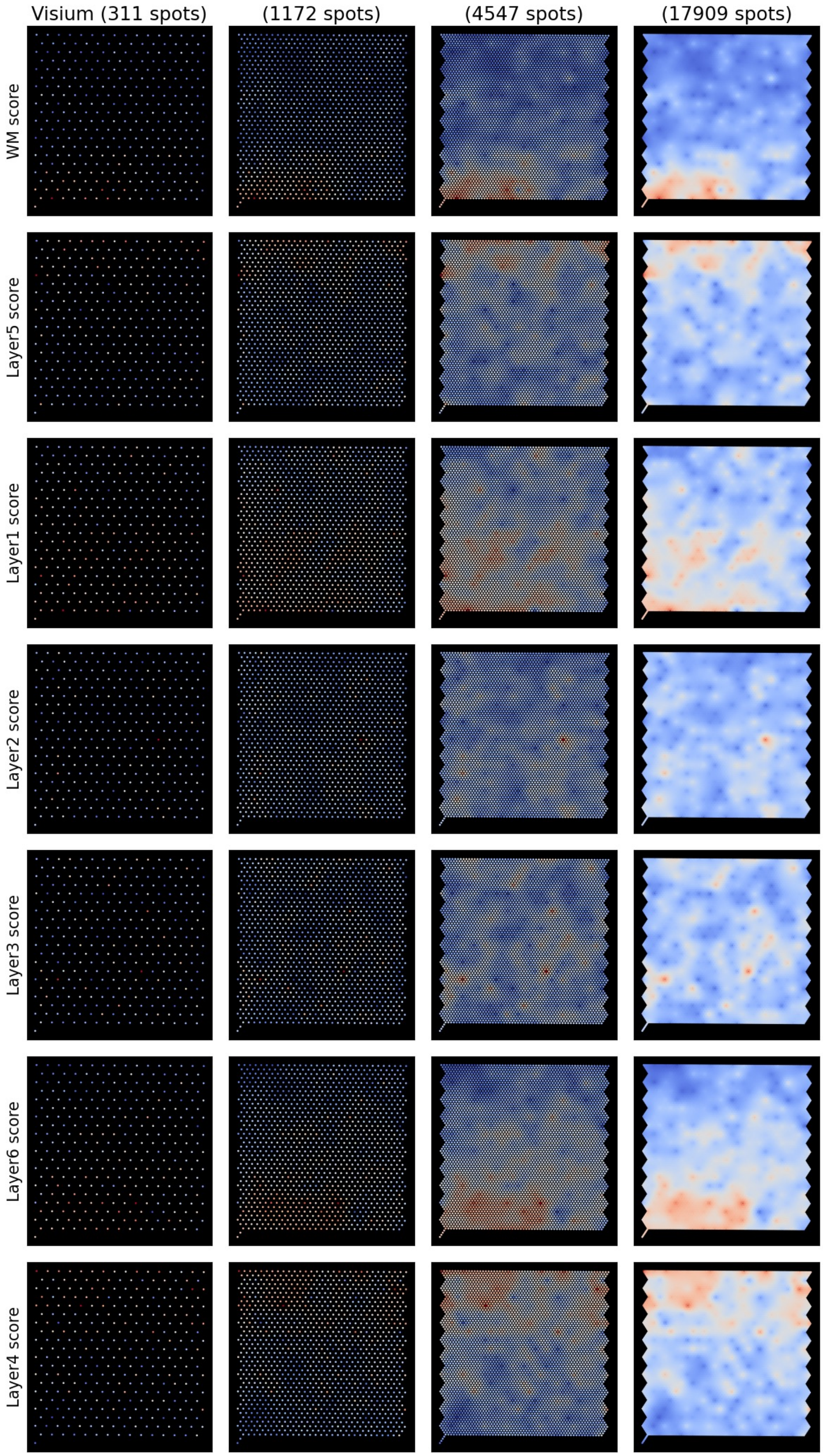


**Supplementary Figure 5: Resolution enhancement of gene signature scores in the DLPFC dataset (151673).**
The figure depicts the enhancing of the spatial resolution of gene signature scores in the DLPFC slide 151673. The 10x Visium original slide was cropped (first column) to visualize the enhancement at different resolutions. Gene signature scores were defined per region using 'scanpy.tl.score_genes' as the weighted scaled average of top20 marker genes of that region (adjusted for gene background).

### Supplementary Figure 6

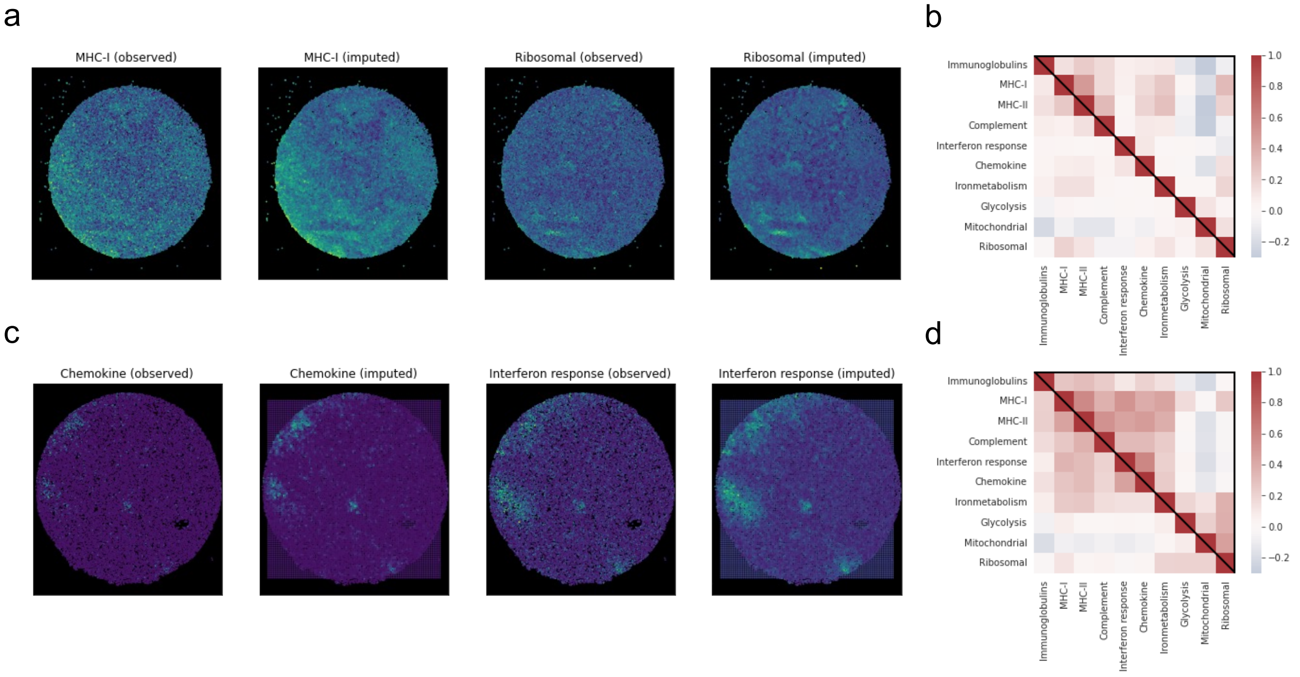


**Supplementary Figure 6: Application of Smoother for smoothing and enhancing functional activity scores in the Slide-seqV2 MBM data.**

The figure demonstrates the imputation (a, b) and resolution enhancement (c, d) of functional signature scores in a Slide-seqV2 melanoma brain metastasis slide (MBM11_rep2, a, b) and an extracranial melanoma metastasis slide (ECM01_rep2, c, d). Functional scores are calculated based on marker gene expression following the original publication. (a) MHC-I and ribosomal scores before and after smoothing. (b) Pairwise correlation between functional signature scores before (lower left) and after (upper right) smoothing. (c) Resolution enhancement of chemokine and interferon response scores before and after enhancement. (d) Pairwise correlation between functional signature scores before (lower left) and after (upper right) enhancement.

Supplementary Figure 7

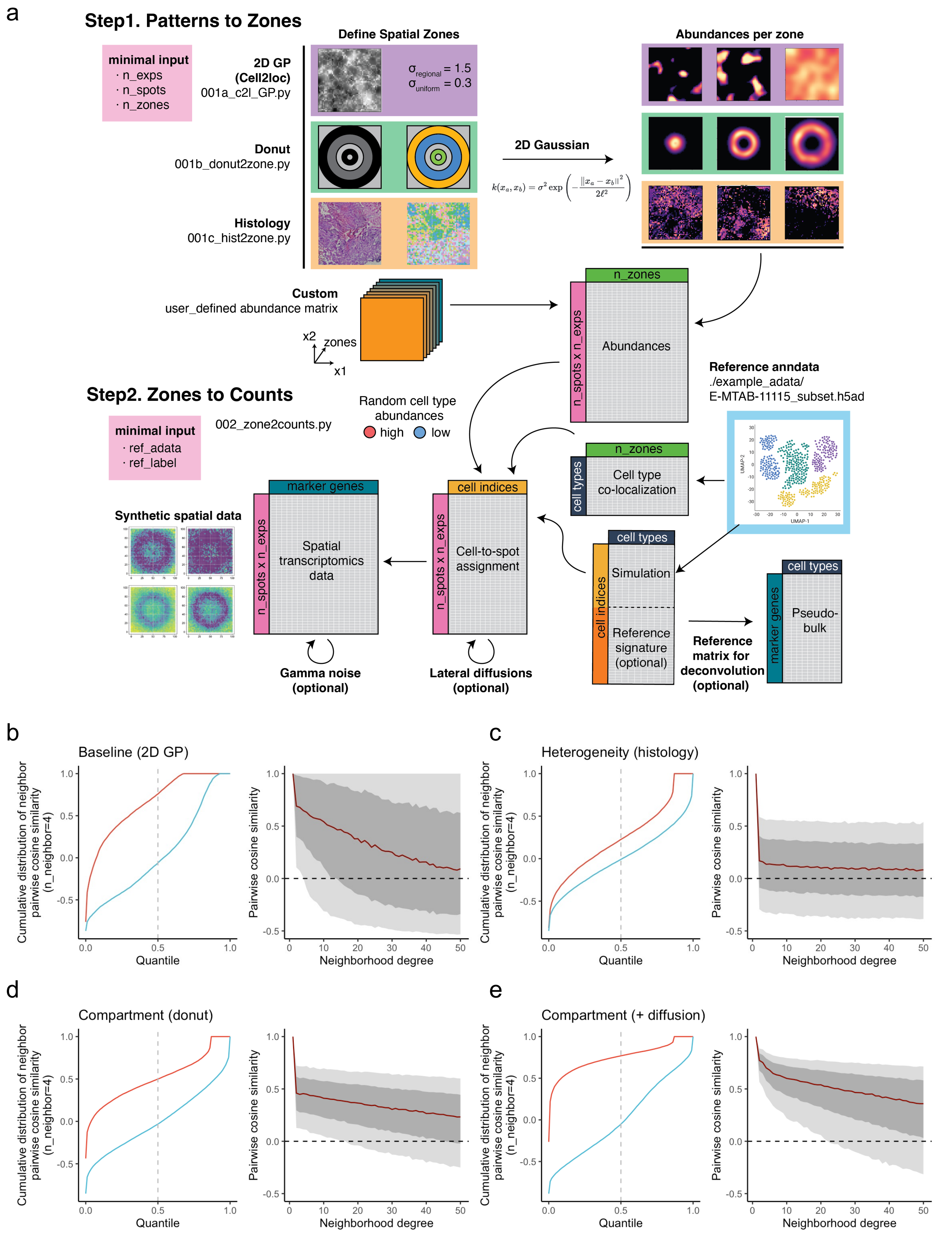


**Supplementary Figure 7: Simulating spatial transcriptomics data for deconvolution benchmark.**
(a) The simulation pipeline, based on cell2location(10) introduces key modifications to assign arbitrary spatial patterns to zones (co-localized cell type groups) and incorporate additional noise (termed lateral diffusion in this paper, also called contamination, spot swapping or bleeding in literature) to account for cell-type-independent dependencies. (b-e) Spatial dependency structures of the simulated data, related to **Supplementary Figure 1**. From b to e: a baseline pattern generated by the 2D Gaussian process (cell2location), a heterogeneous pattern specified by a tumor histology image, a pattern with clear doughnut-shaped compartments, and with additional lateral diffusion noise.

Supplementary Figure 8

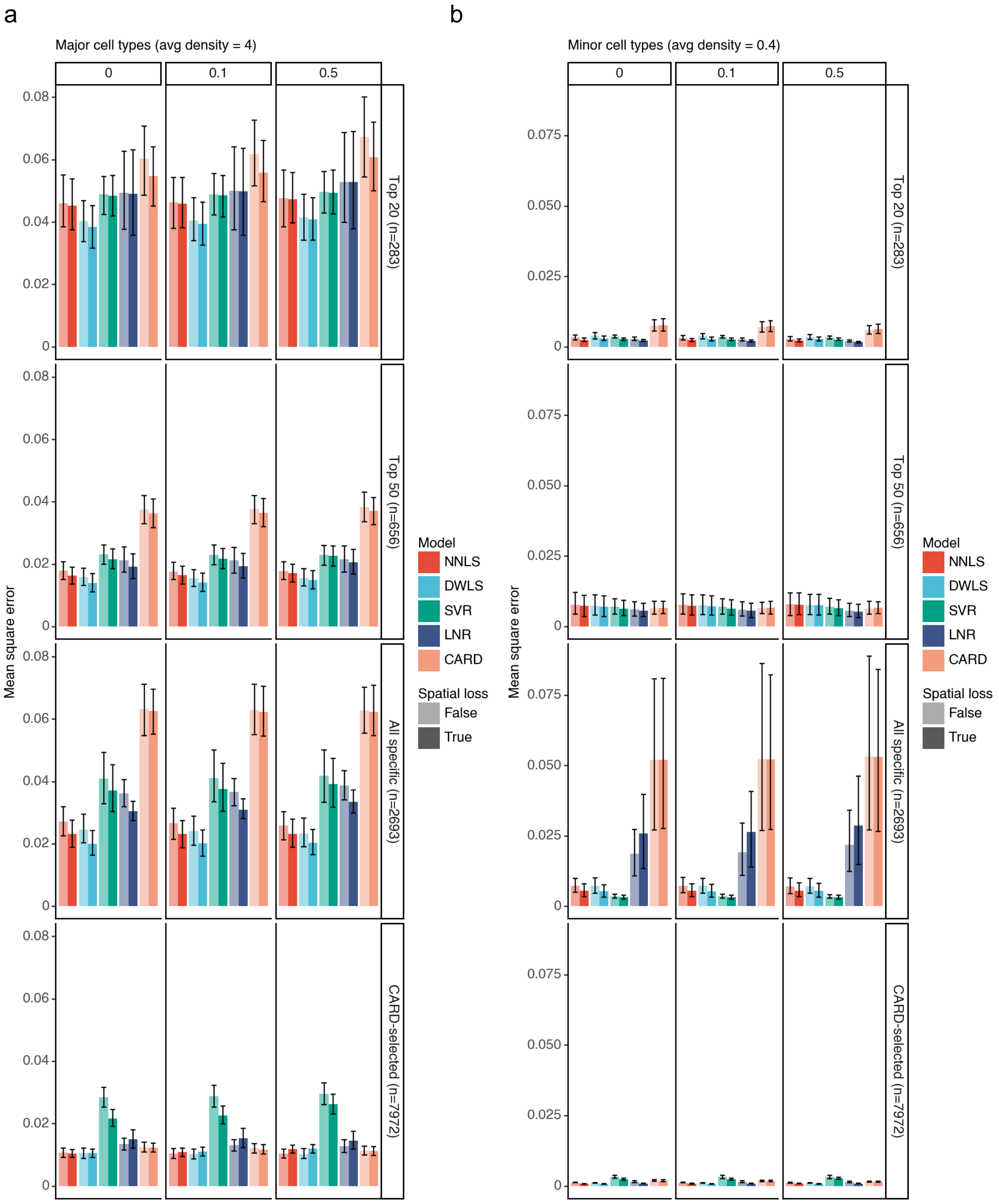


**Supplementary Figure 8: Evaluation of deconvolution accuracy on simulated data under different scenarios measured by mean square error (the lower the better), related to Figure 3a-d.**

Doughnut-shaped spatial transcriptomics datasets, with varying degree of lateral diffusion (columns, number indicates the proportion of mRNA shared with adjacent spots), were generated from scRNA-seq reference on a 50x50 grid. In each of the ten experiments (replicates), we assigned 5 major cell types with high average abundances (a) and 10 minor cell types with low abundances (b) to overlapping spatial compartments. Mean square error (MSE) was calculated based on the true and estimated cell type proportions (sum to one per spot). Each row represents the scenario where only a subset of genes is informative as deconvolution input. From top to bottom: the union of the top 20 marker genes for each cell type (n=283), the union of top 50 marker genes (n=656), all informative markers genes whose log2(fold change) passes the threshold (by default 1) for one and only one cell type (n=2693), and informative genes selected by the CARD model (n=7972). Error bars denote standard error of the mean over 5 high- or 10 low-density cell types in 10 replicates.

### Supplementary Figure 9

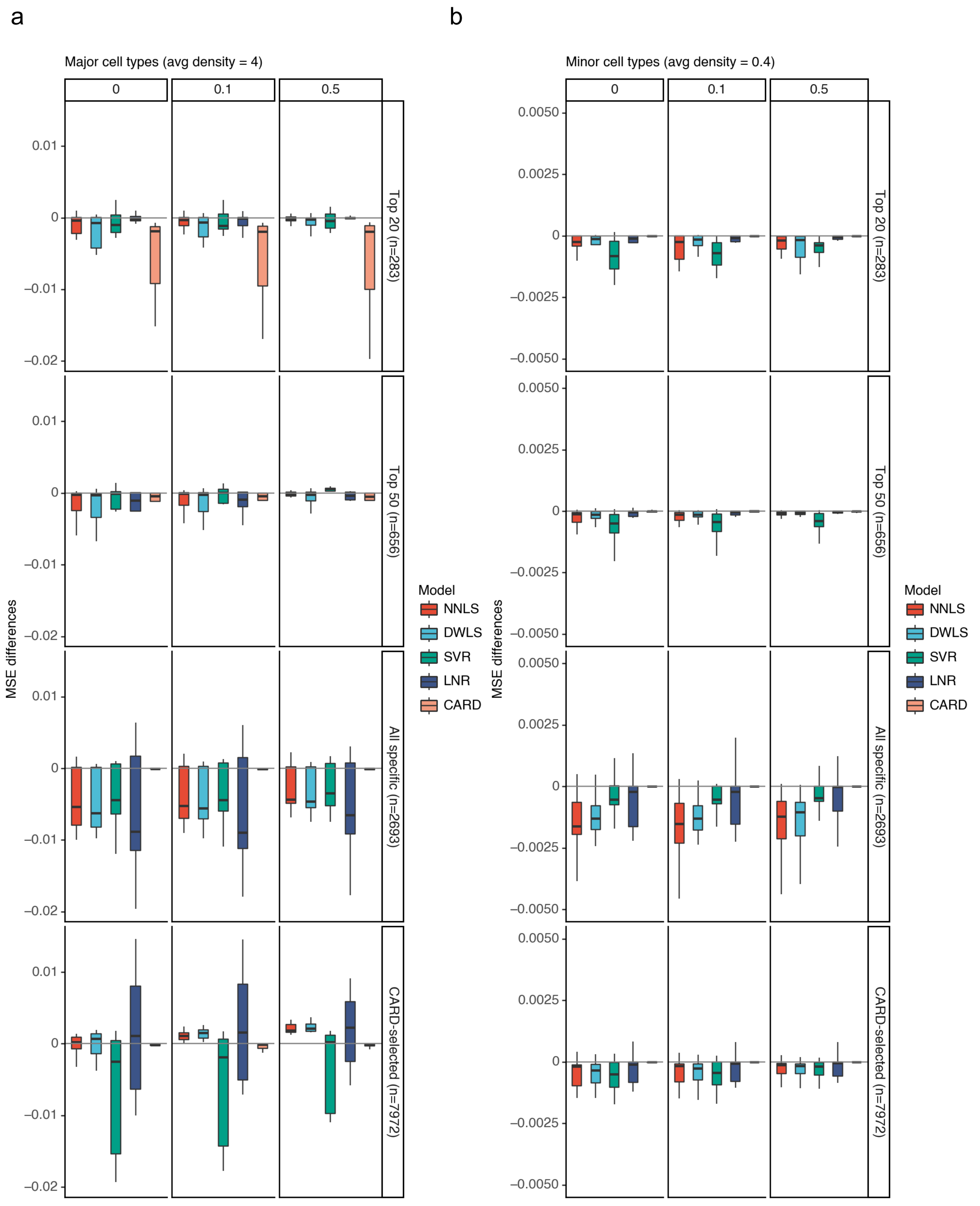


**Supplementary Figure 9: Performance gained from the spatial regularization under different scenarios, measured by mean square error (the lower the better), related to Supplementary Figure 8.**
Box plots showing the distribution of performance differences after the spatial regularization is incorporated into deconvolution. Scores were calculated by subtracting the MSE of the non-spatial baseline model from the MSE of a spatially aware model for each cell type in each experiment. Each box plot demonstrates the median and the 25th/75th percentiles of 50 samples for major cell types (a) and 100 for minor cell types (b), as well as the above and below 1.5 times interquartile ranges indicated by the whiskers.

Supplementary Figure 10

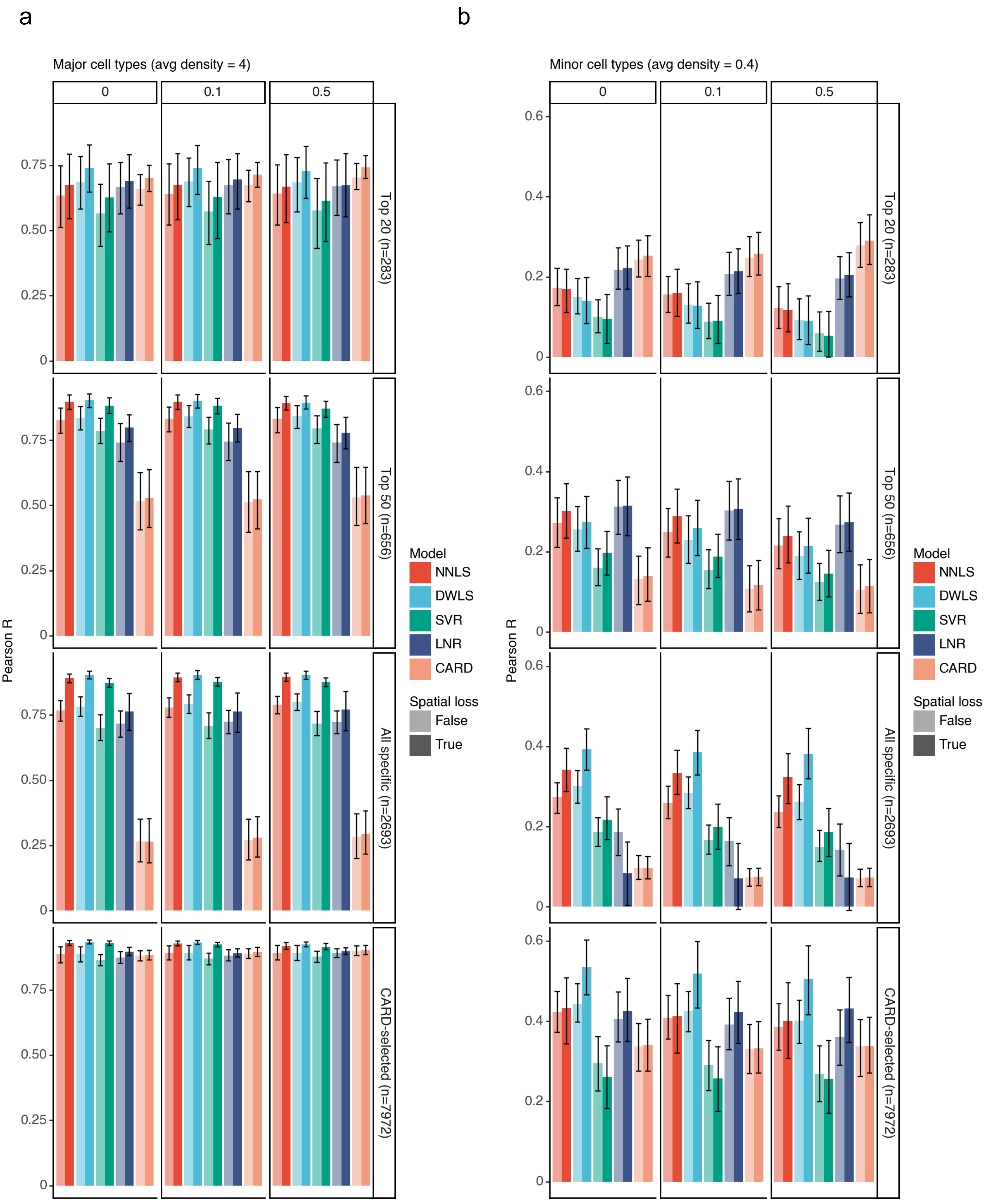


**Supplementary Figure 10: Evaluation of deconvolution accuracy on simulated data under different scenarios measured by Pearson correlation (the higher the better), related to Figure 3a-d.**
Similar to **Supplementary Figure 8** except the performance is measured by the Pearson correlation between the true and estimated cell-type proportions (sum to one per spot).

Supplementary Figure 11

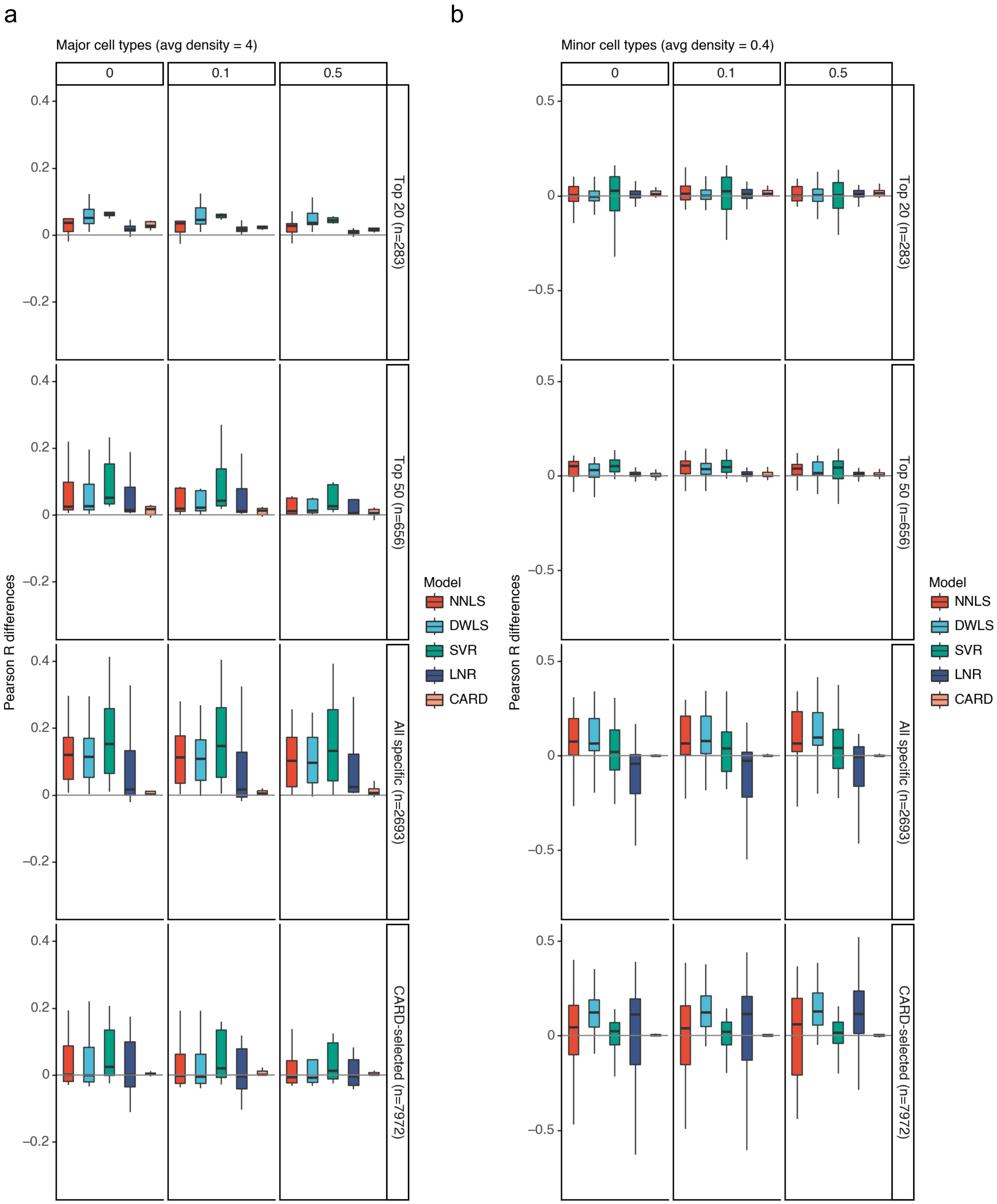


**Supplementary Figure 11: Performance gained from the spatial regularization under different scenarios, measured by Pearson correlation (the higher the better), related to Supplementary Figure 10.**
Similar to **Supplementary Figure 9** except the performance is measured by the Pearson correlation between the true and estimated cell-type proportions (sum to one per spot).

### Supplementary Figure 12

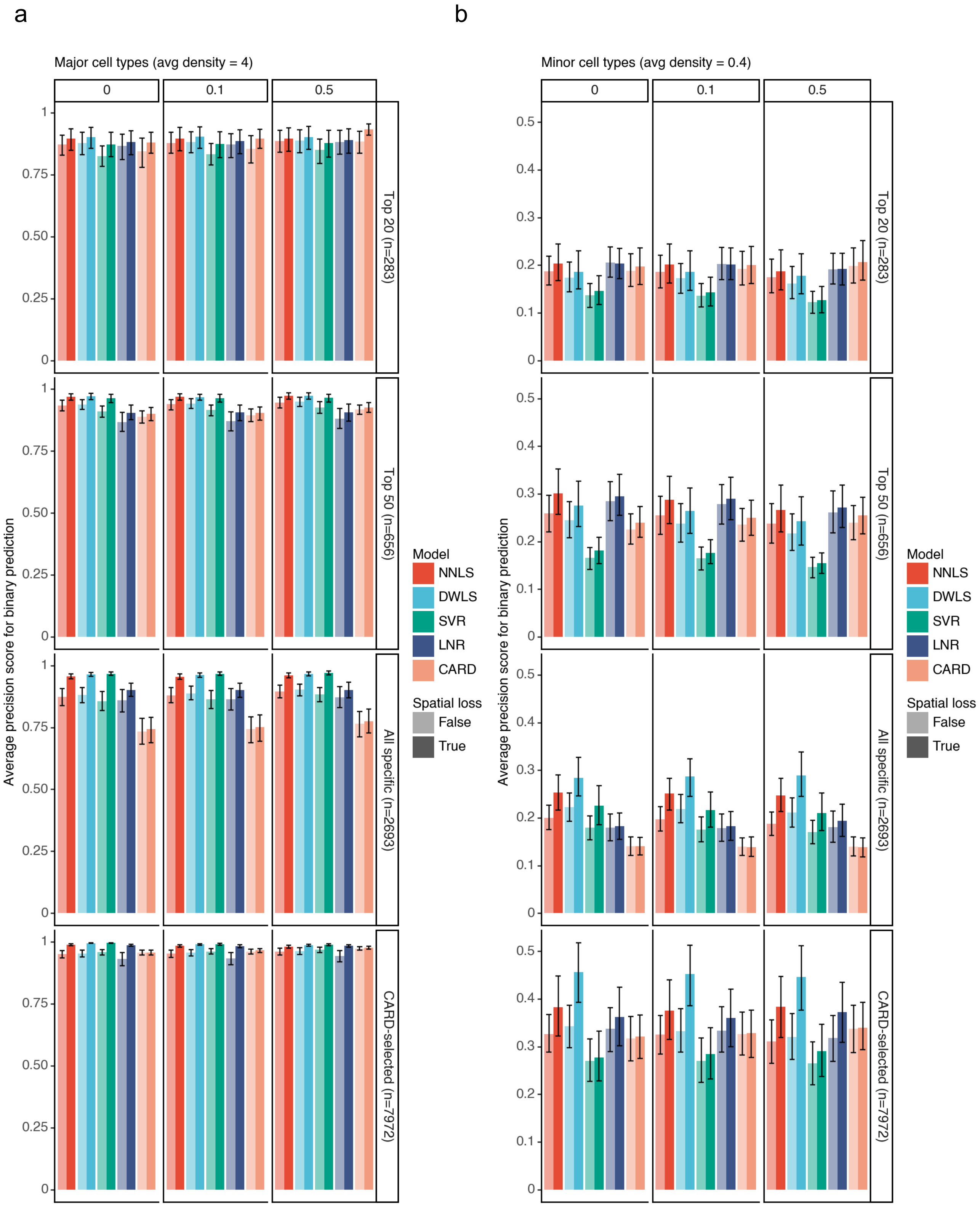


**Supplementary Figure 12: Evaluation of deconvolution accuracy on simulated data under different scenarios measured by binary prediction accuracy (the higher the better), related to Figure 3a-d.**
Similar to **Supplementary Figure 8** except the performance is measured by binary prediction accuracy. Cell types that have a ground truth proportion larger than 0.05 were considered as present at a spot. Average precision scores were calculated for cell types that are at least present in one location in each experiment individually.

Supplementary Figure 13

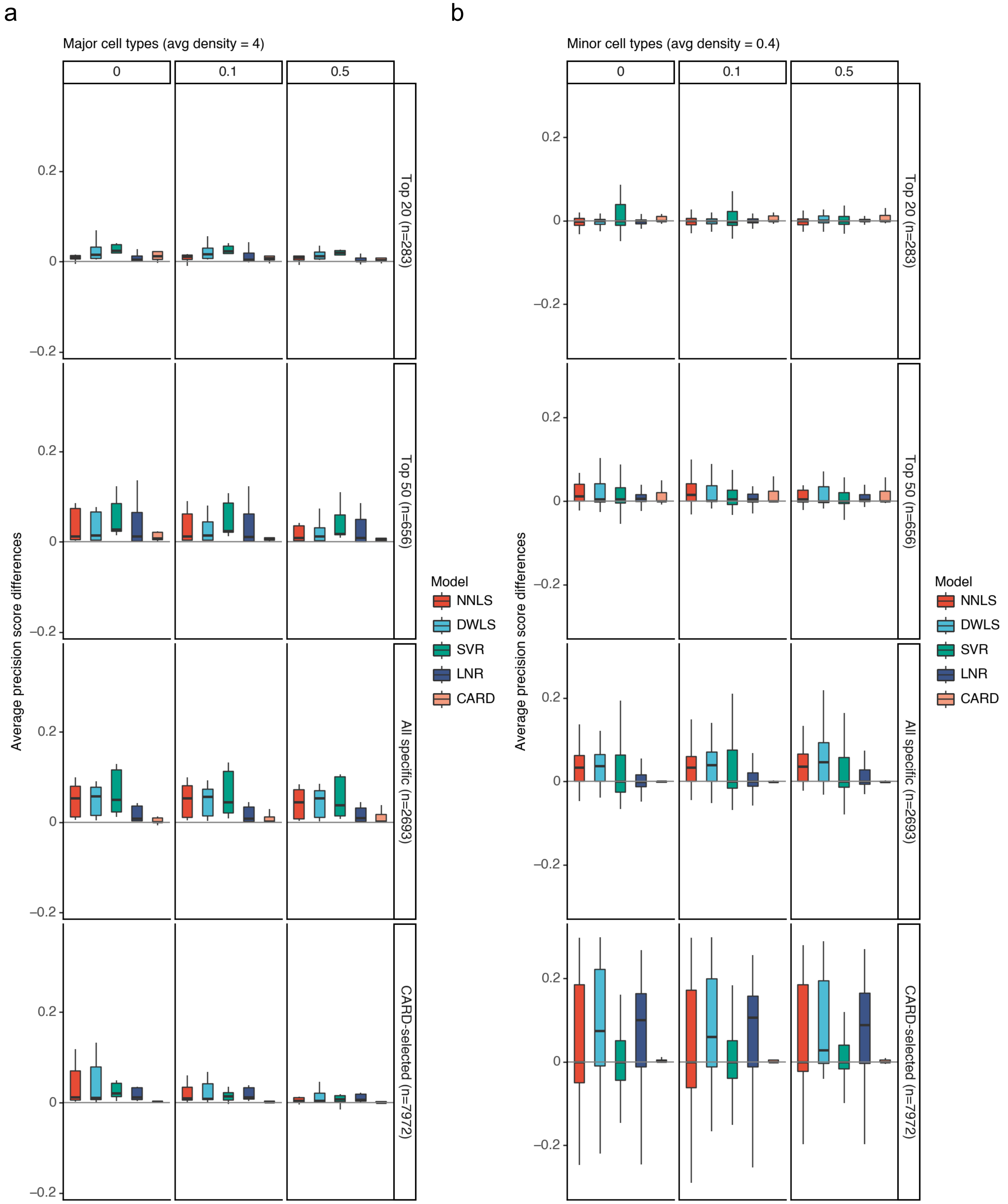


**Supplementary Figure 13: Performance gained from the spatial regularization under different scenarios, measured by binary prediction accuracy (the higher the better), related to Supplementary Figure 12.**
Similar to **Supplementary Figure 9** except the performance is measured by the average precision score.

Supplementary Figure 14

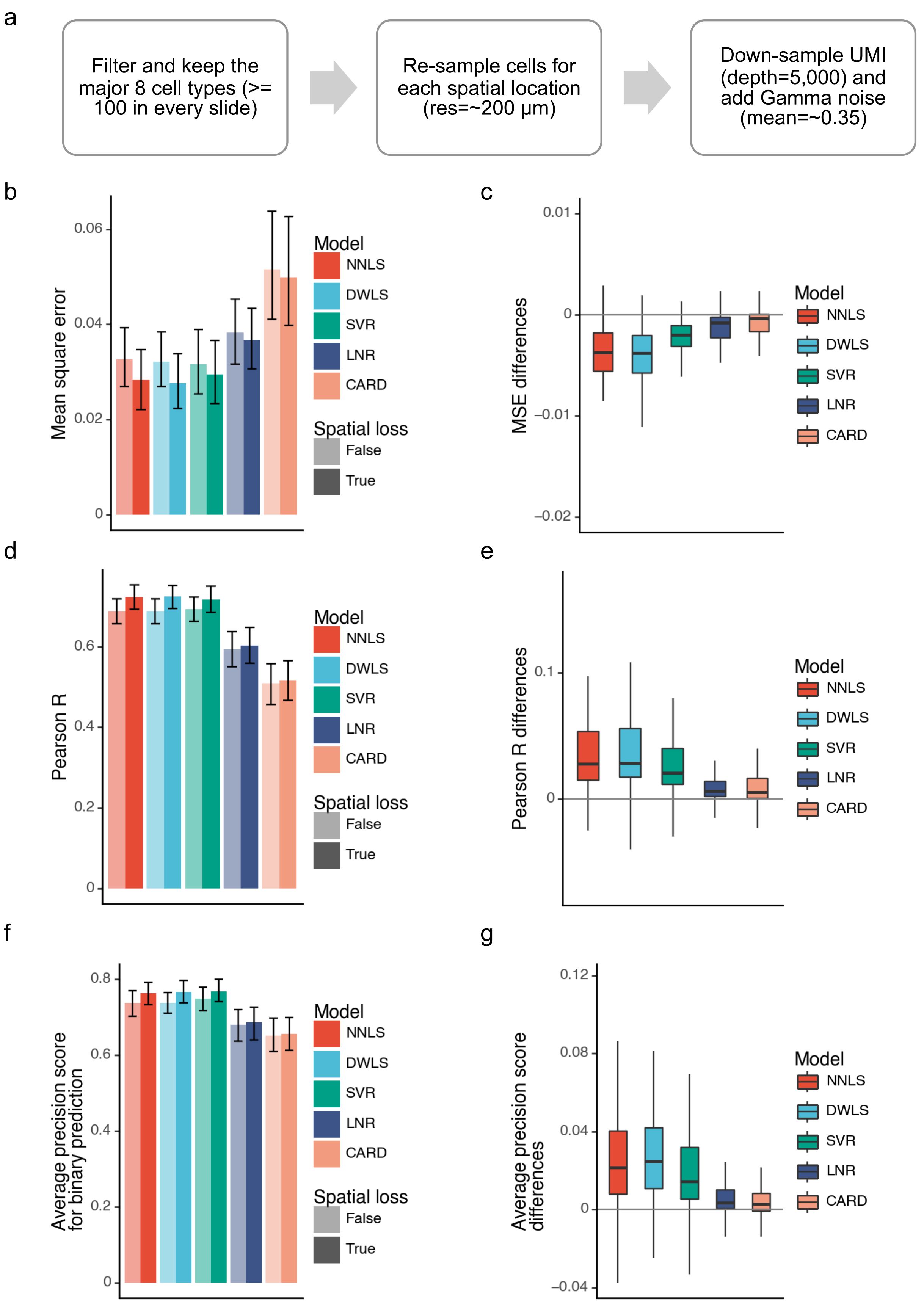


**Supplementary Figure 14: Evaluation of deconvolution performance on the Sci-Space data of mouse embryo, related to Figure 3e-h.**

(a) Schematic overview of the data simulation pipeline. (b-g) Deconvolution performance as measured by mean square error (b, c), Pearson correlation (d, e), and average precision score for binary prediction (f, g). Error bars denote standard error of the mean over all 8 cell types across 14 slides (112 data points in total). Box plots show median, 25th/75th percentiles of the 112 samples, and whiskers indicating 1.5 times the interquartile ranges.

Supplementary Figure 15

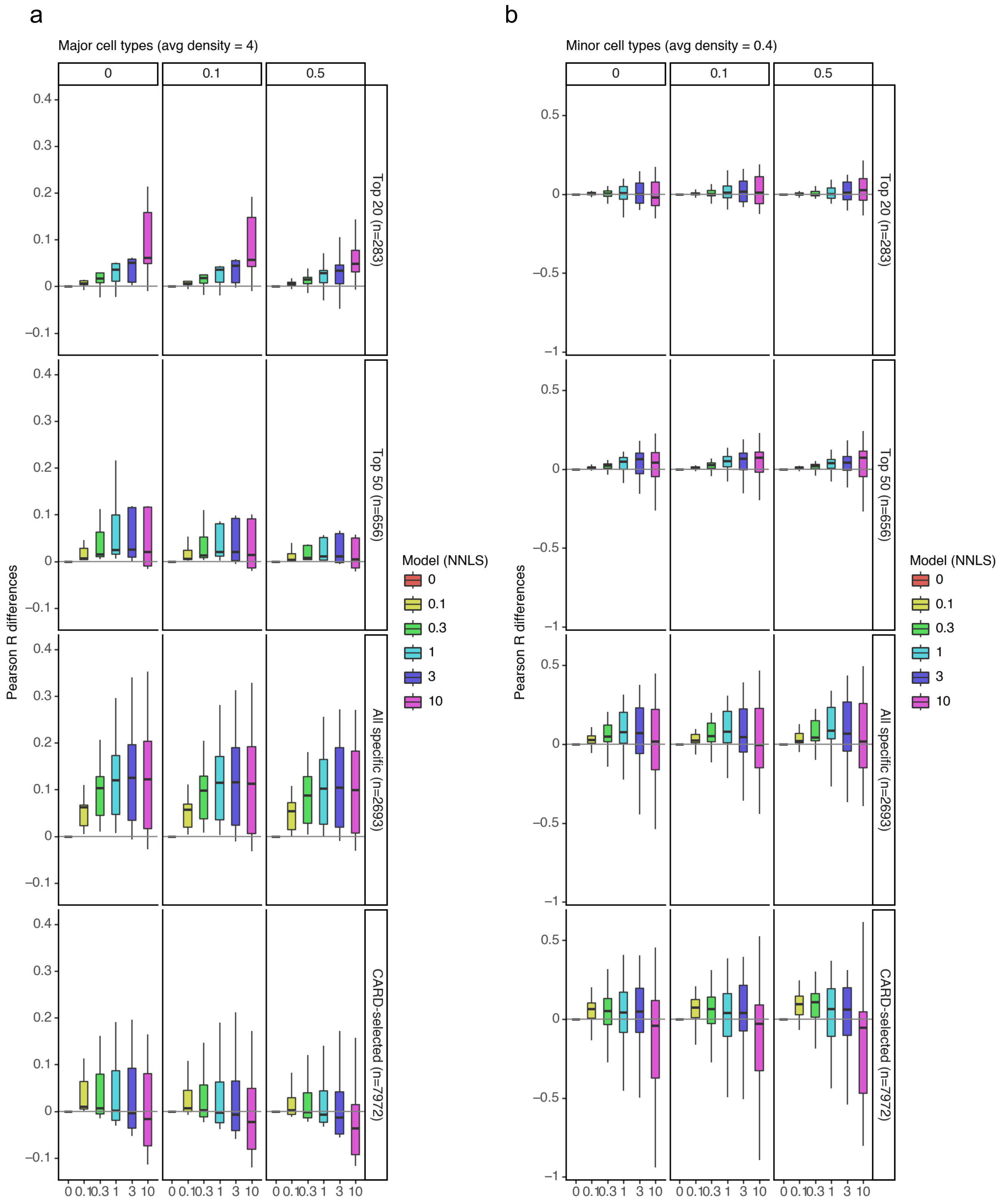


**Supplementary Figure 15: Robust beneficial effect of spatial regularization in NNLS-based deconvolution.**
Box plots display the distribution of deconvolution performance differences after incorporating spatial losses with varying strengths (λ_sp, x-axis). Performance was measured by Pearson correlation, and the coefficient of a non-spatial baseline NNLS model was subtracted from the value of the spatially aware NNLS with different λ_sp for each cell type in each experiment. Each box plot demonstrates the median and the 25th/75th percentiles, and whiskers indicating 1.5 times interquartile ranges indicated by the whiskers of 50 samples for major cell types (a) and 100 for minor cell types (b).

Supplementary Figure 16

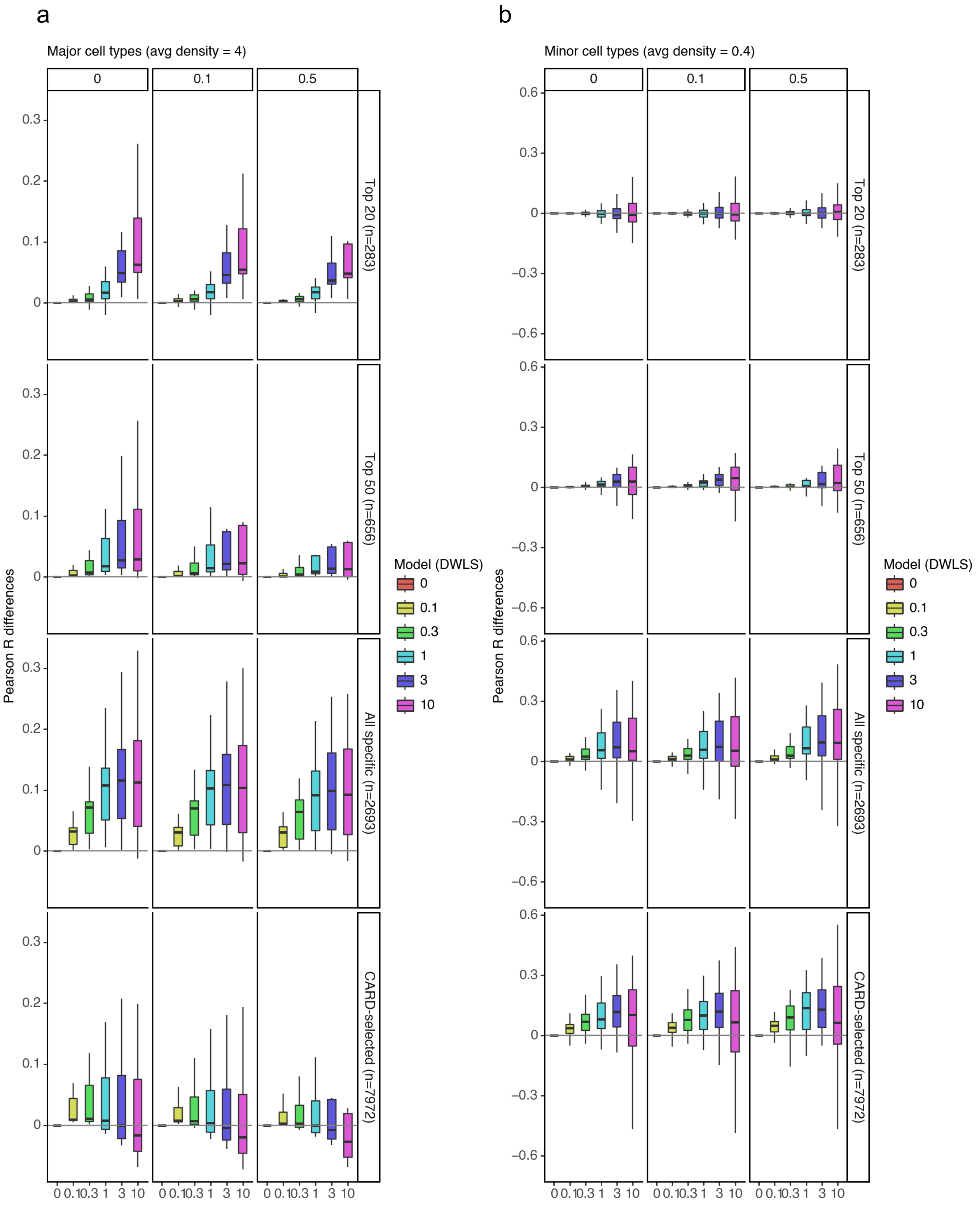


**Supplementary Figure 16: Robust beneficial effect of spatial regularization in DWLS-based deconvolution.** Similar to **Supplementary Figure 15** except the base non-spatial deconvolution model is DWLS.

Supplementary Figure 17

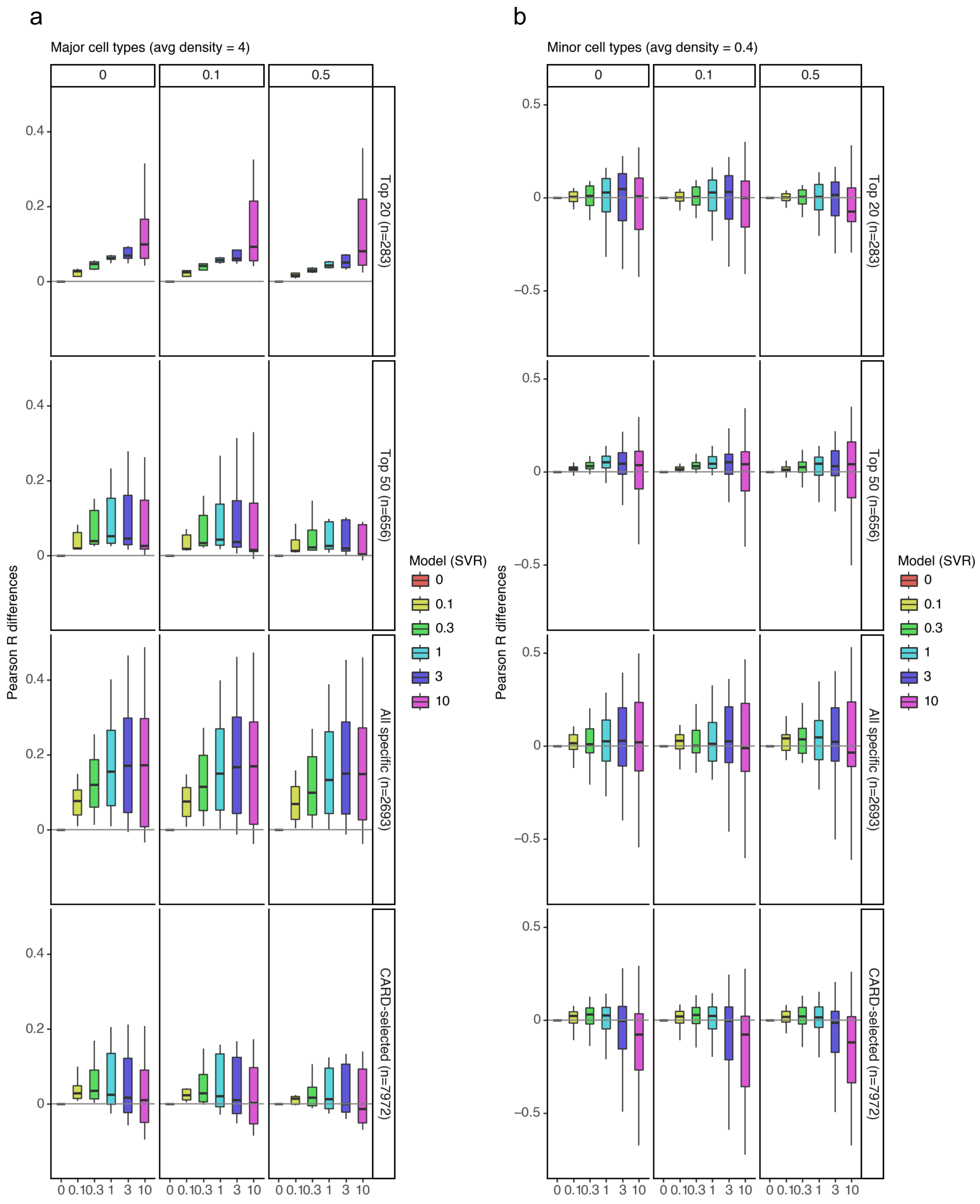


**Supplementary Figure 17: Robust beneficial effect of spatial regularization in SVR-based deconvolution.**
Similar to **Supplementary Figure 15** except the base non-spatial deconvolution model is nu-SVR.

Supplementary Figure 18

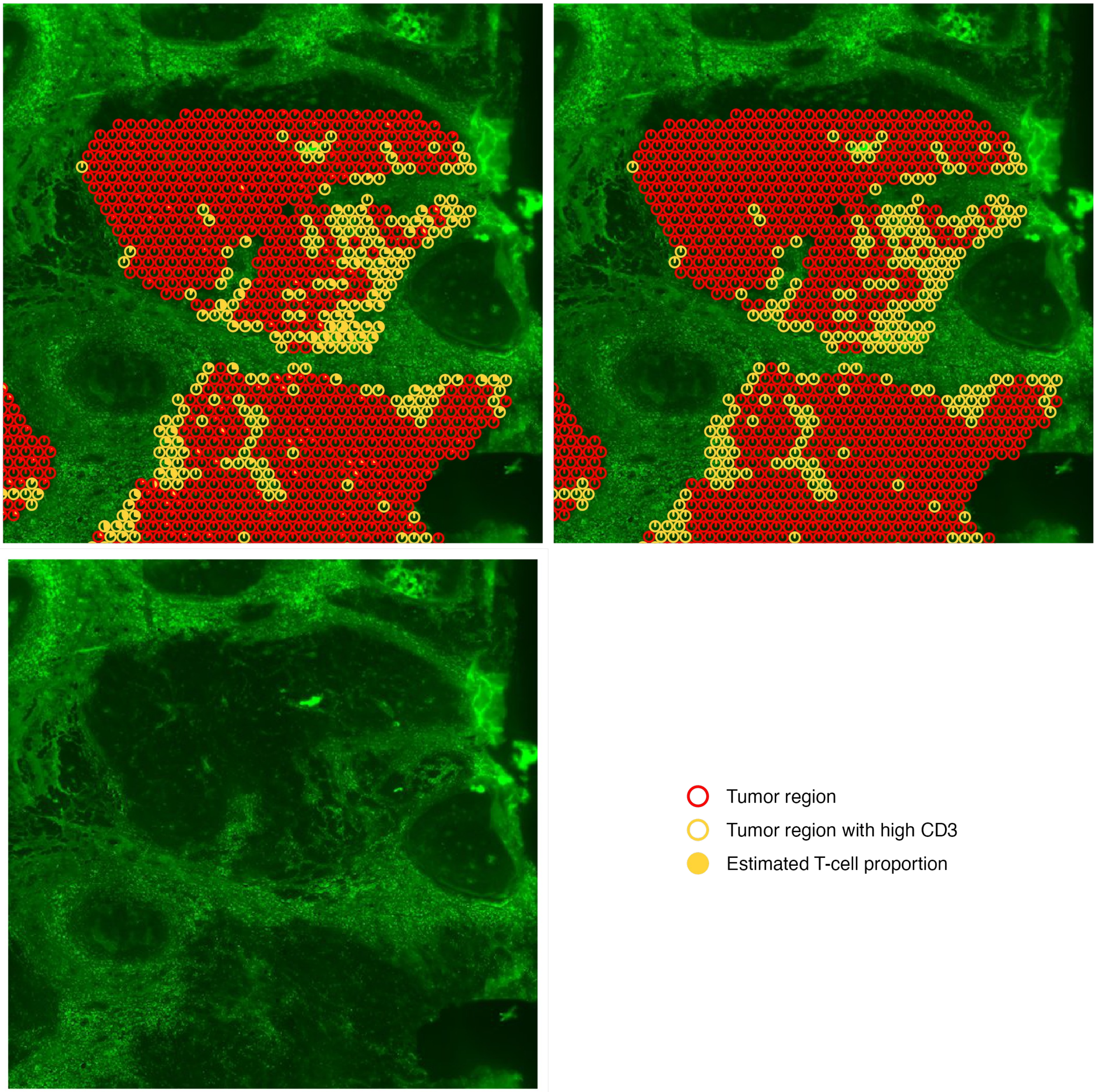


**Supplementary Figure 18: T-cell infiltration in the zoom-in tumor region of the ductal carcinoma section, related to Figure 4a-c.**
Each circle denotes a tumor occupied spot based on pathological annotation. Spots were classified as CD3+ (yellow) or CD3- (red) according to a two-component Gaussian mixture model on CD3 intensity. Pie charts indicate the estimated T-cell proportions at each spot. Background shows the single channel CD3 immunofluorescence staining (FITC/green) intensity.

Supplementary Figure 19

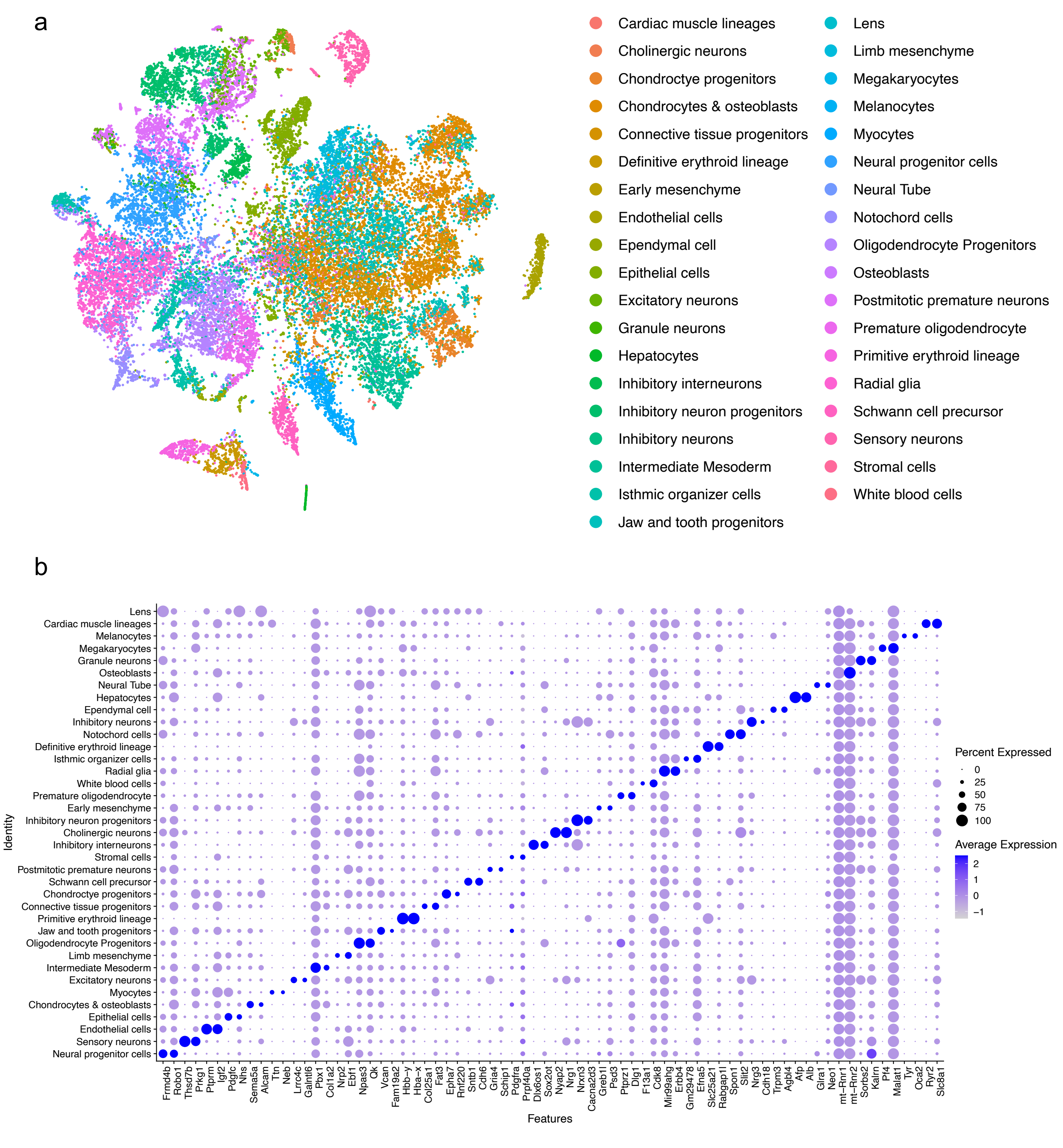


**Supplementary Figure 19: T-SNE visualization of and marker expression of the scRNA-seq reference from the Mouse Organogenesis Cell Atlas (MOCA), related to Figure 4h-k.**

Supplementary Figure 20

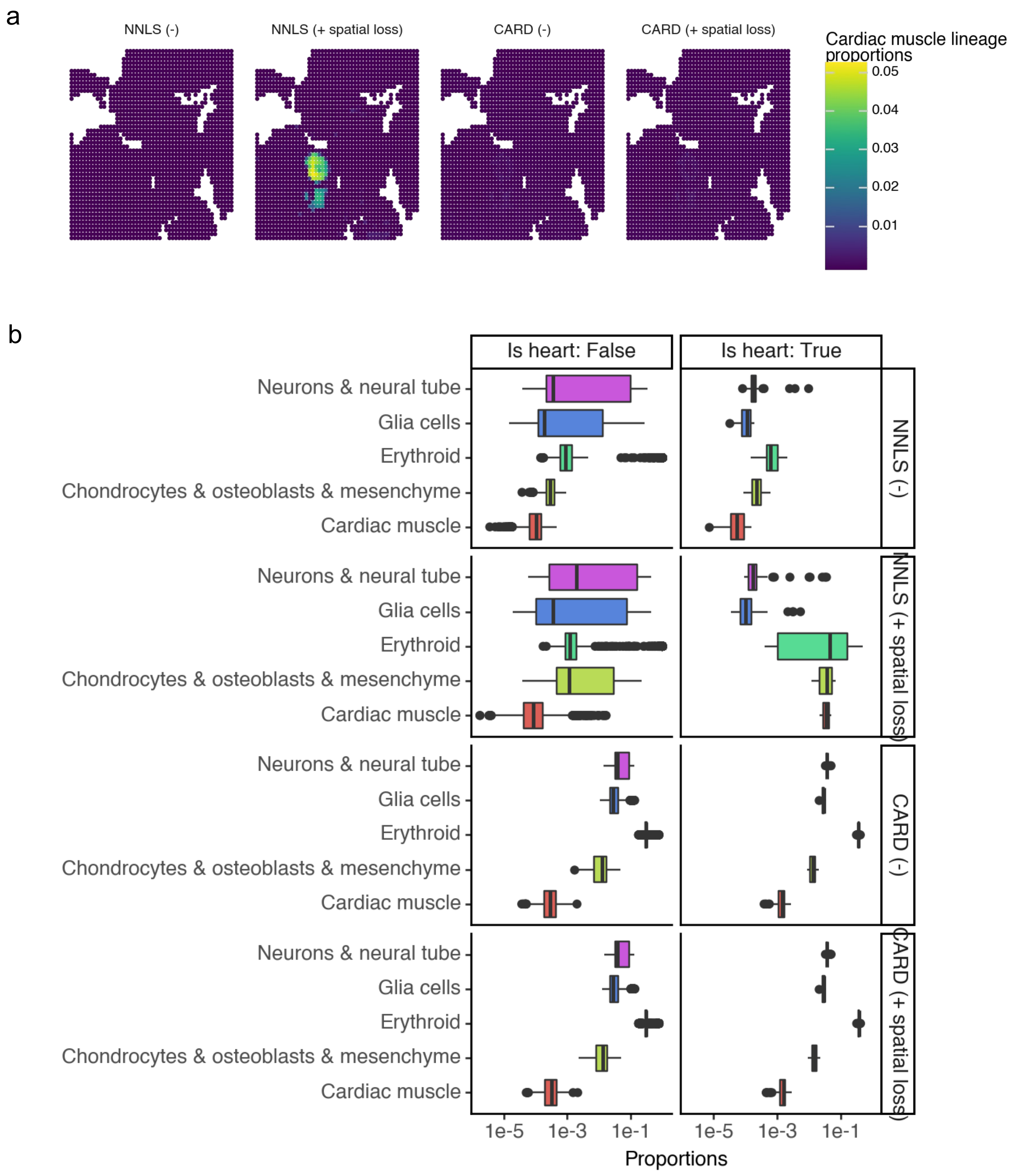


**Supplementary Figure 20: Detailed examination of deconvolution of cardiac muscle lineage in the spatial-CUT&Tag dataset, related to Figure 4h-k.**

(a) Estimated proportions of cardiac muscle lineage. (b) Distributions of estimated proportions of the five major lineages in and outside the heart region. The heart region is defined by NNLS (+ spatial loss)-estimated proportion >= 0.02. List of cell types in each lineage: ‘Neurons & neural tube’: cholinergic neurons, excitatory neurons, granule neurons, neural progenitor cells, neural tube, notochord cells, sensory neurons, inhibitory interneurons, inhibitory neuron progenitors, postmitotic premature neurons, inhibitory neurons. ‘Glia cells’: oligodendrocyte progenitors, premature oligodendrocyte, ependymal cell, Schwann cell precursor, radial glia. ‘Erythroid’: primitive erythroid lineage, definitive erythroid lineage, megakaryocytes, white blood cells. ‘Chondrocytes & osteoblasts & mesenchyme’: chondrocyte progenitors, chondrocytes & osteoblasts, osteoblasts, connective tissue progenitors, early mesenchyme, limb mesenchyme'. ‘Cardiac muscle’: cardiac muscle lineages, myocytes.

### Supplementary Figure 21

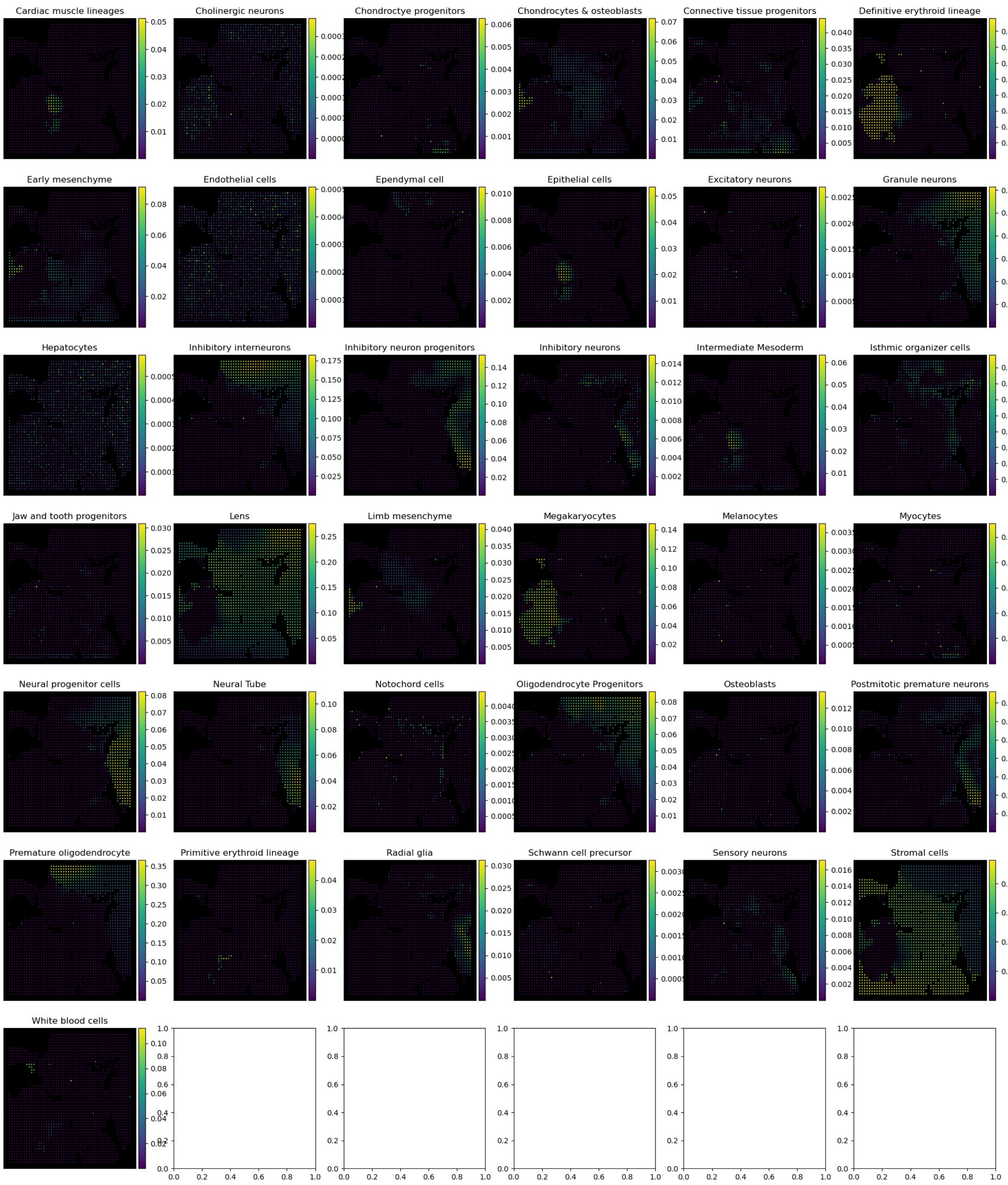


**Supplementary Figure 21: Per-cell-type deconvolution results of NNLS (+ spatial loss) in the spatial-CUT&Tag dataset, related to Figure 4h-k.**

### Supplementary Figure 22

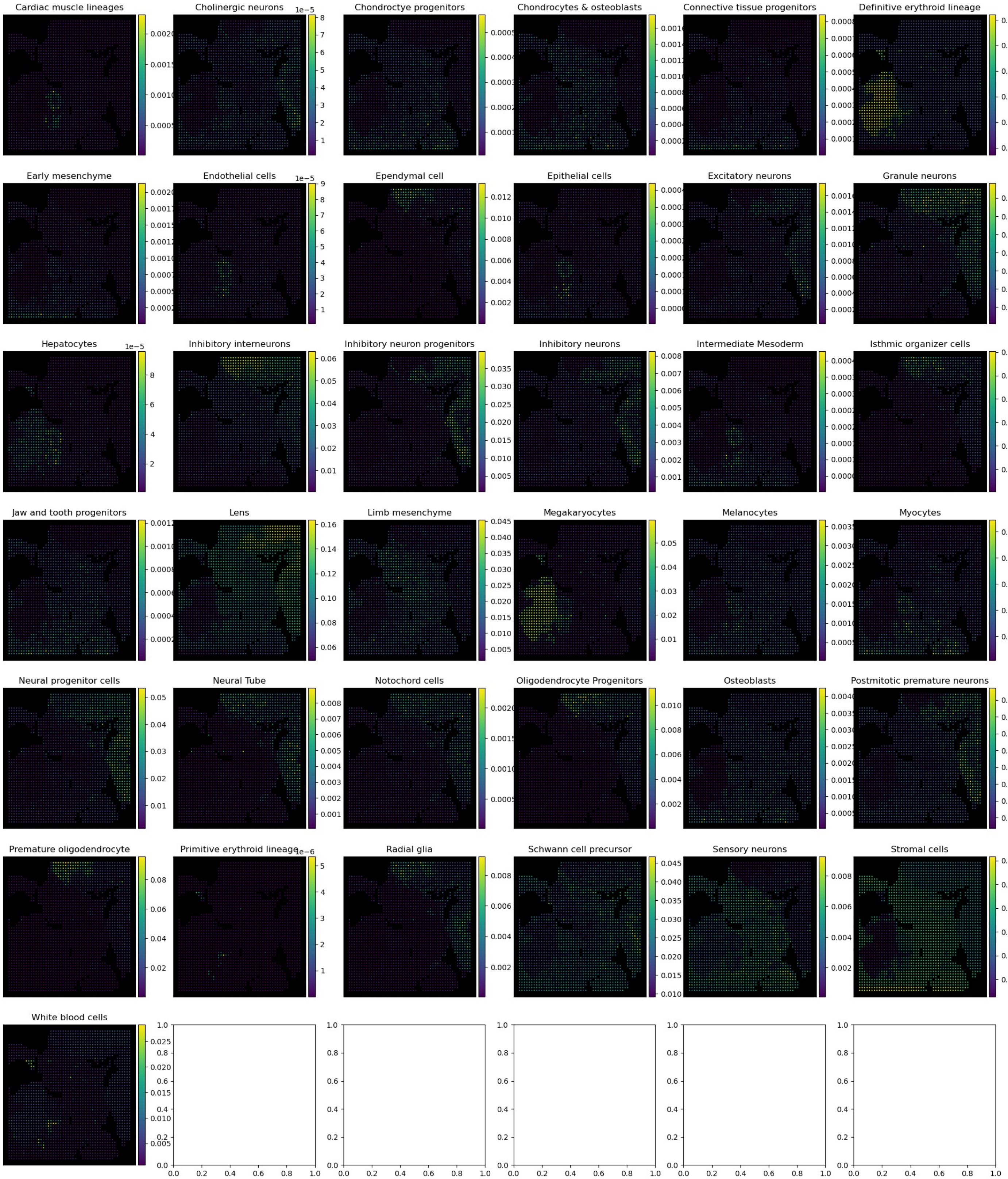


**Supplementary Figure 22: Per-cell-type deconvolution results of CARD (phi=0.99) in the spatial-CUT&Tag dataset, related to Figure 4h-k.**

Supplementary Figure 23

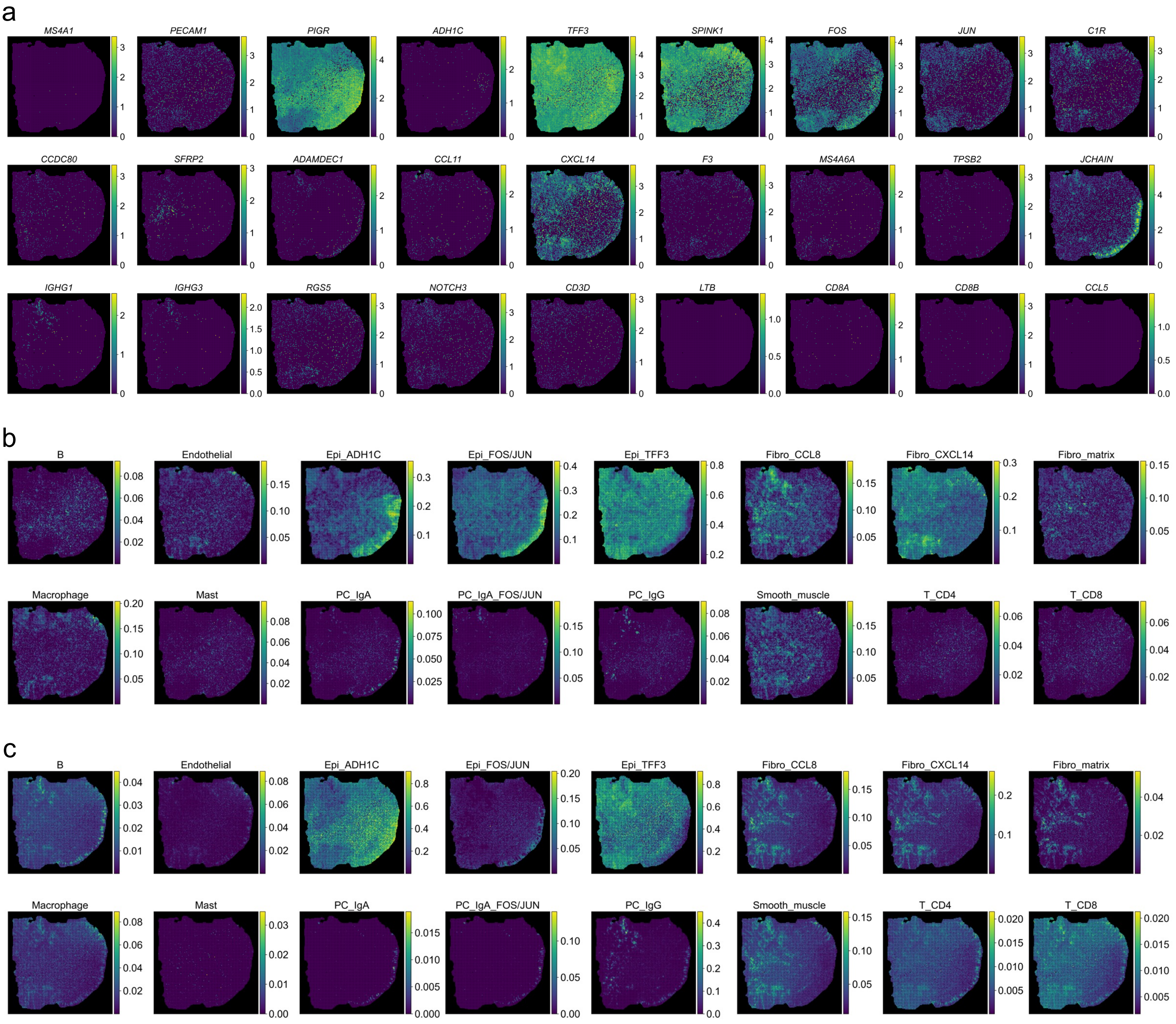


**Supplementary Figure 23: Full deconvolution results and marker gene expression in the CRC tumor tissue Stereo-seq section, related to Figure 5.**
(a) Spatial expression of cell-type specific marker genes. (b-c) Cell type proportions as predicted by Smoother-guided DWLS (b) and CARD with spatial regularization (c).

### Supplementary Figure 24

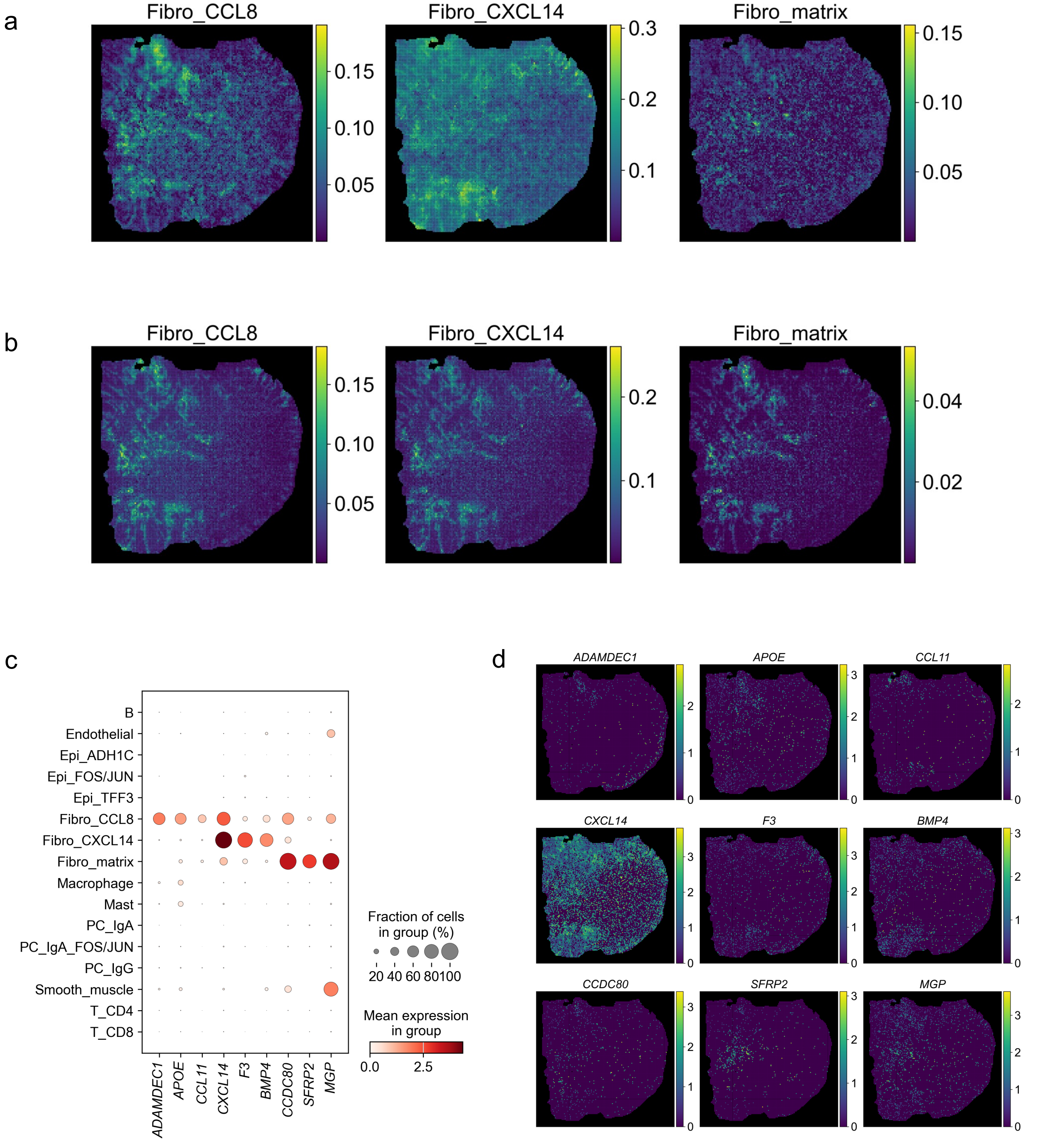


**Supplementary Figure 24: Comparison between fibroblast subtype deconvolution results and marker gene expression in the CRC tumor tissue Stereo-seq section, related to Figure 5.**
(a-b) Fibroblast subtype proportions as predicted by Smoother-guided DWLS (a) and CARD with spatial regularization (b). (c) Expression dotplot of three marker genes for each fibroblast subtype in the reference scRNA-seq samples, which correspond to previously derived ADAMDEC1+/CCL8+, CXCL14+ and matrix transcriptional programs (Pelka et al.(11)). (d) Spatial expression of the subtype-specific markers shown in (c).

### Supplementary Figure 25

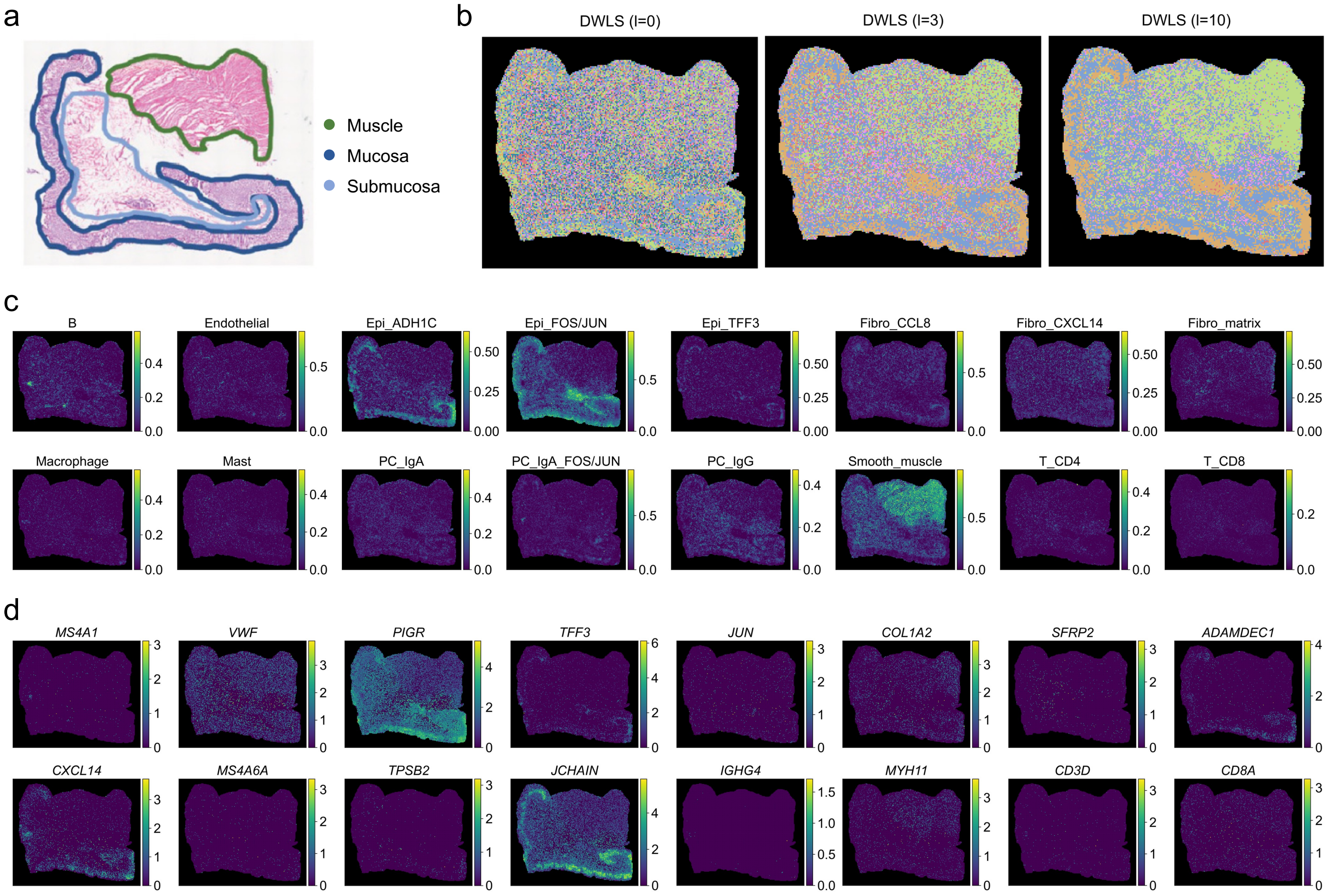


**Supplementary Figure 25: Deconvolution results and marker gene expression in the adjacent healthy tissue Stereo-seq section of the same CRC patient, related to Figure 5.**
(a) Pathology annotation of the histology slide, adapted from(8). (b-d) Deconvolution performed using the same reference expression matrix as the tumor section. (b) Slide subregions identified by clustering based on Smoother-estimated cell-type compositions, with increasing spatial loss from left to right. (c) Cell type proportions as predicted by Smoother-guided DWLS with spatial loss (l=10). (d) Expression of cell-type specific marker genes.

### Supplementary Figure 26

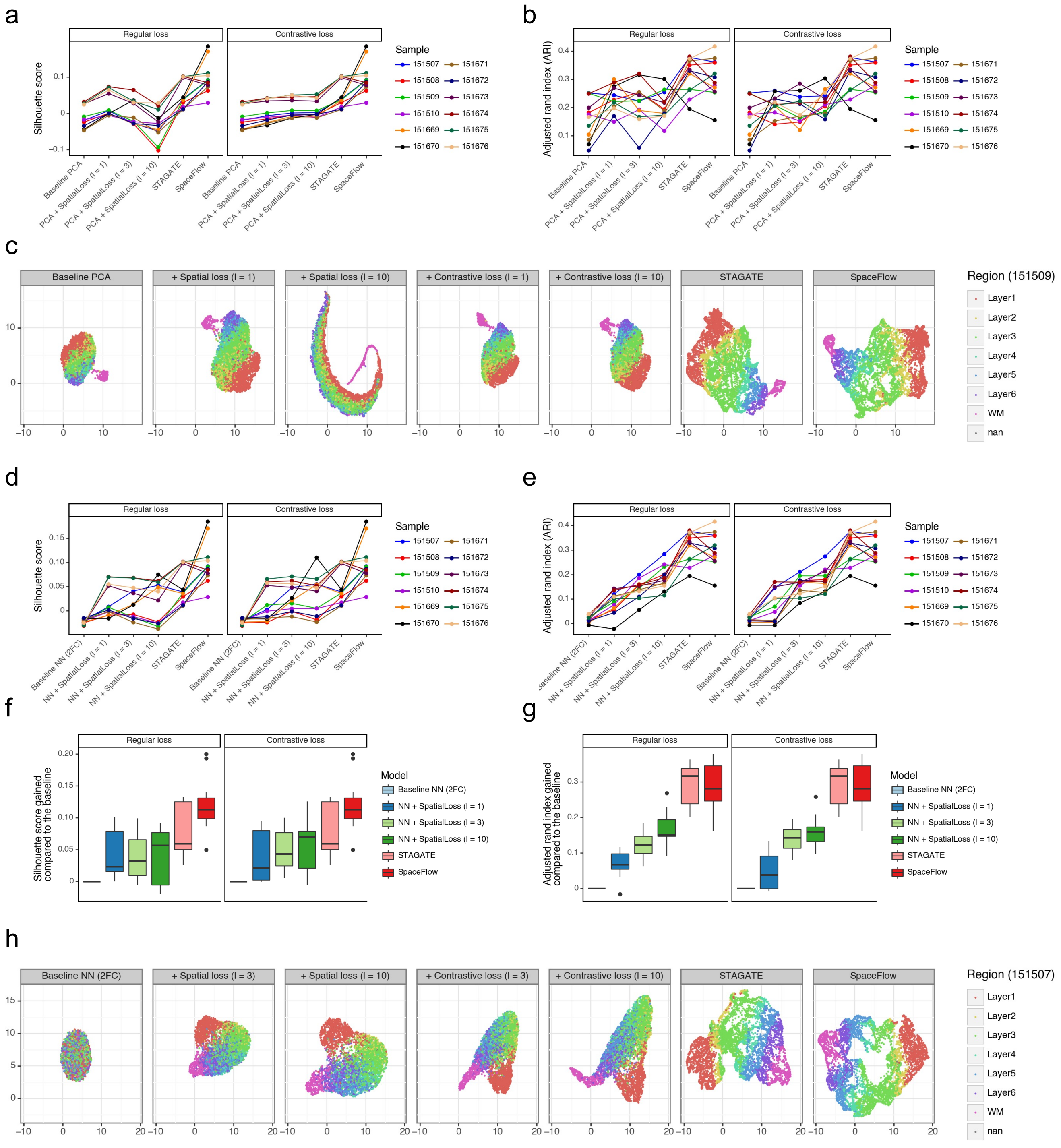


**Supplementary Figure 26: Comparative analyses of dimensionality reduction performance on the DLPFC dataset.**
For each sample, we projected either log-normalized expression (PCA) or the raw counts (other models) of the top 2000 highly variable genes onto a 30-dimension latent space using each model to calculate the Silhouette score. 'Mclust' was used to cluster spots based on the latent embeddings into the same number of clusters (cortex layers) as observed in the sample. Adjusted rand index (ARI) was calculated based on the clustering results. (a-c) Performance comparisons of PCA models with varying strength of the regular spatial loss and the contrastive spatial loss and the two graph neural networks, STAGATE and SpaceFlow. (c) UMAP visualization of the latent representation learned by each model, colored by spot layer membership. (d-h) Performance comparisons of the baseline neural network with varying strength of the regular spatial loss and the contrastive spatial loss and the two graph neural networks, STAGATE and SpaceFlow. The baseline model was constructed by replacing the two graph attention layers in STAGATE with two fully connected layers of the same number of hidden units (128). (f-g) Performance gains were computed against the baseline for each slide. Each boxplot shows the median and the 25th/75th percentiles over the 12 samples and whiskers indicating the 1.5 times interquartile ranges. (h) UMAP visualization of the latent representation learned by each model, colored by spot layer membership.

Supplementary Figure 27

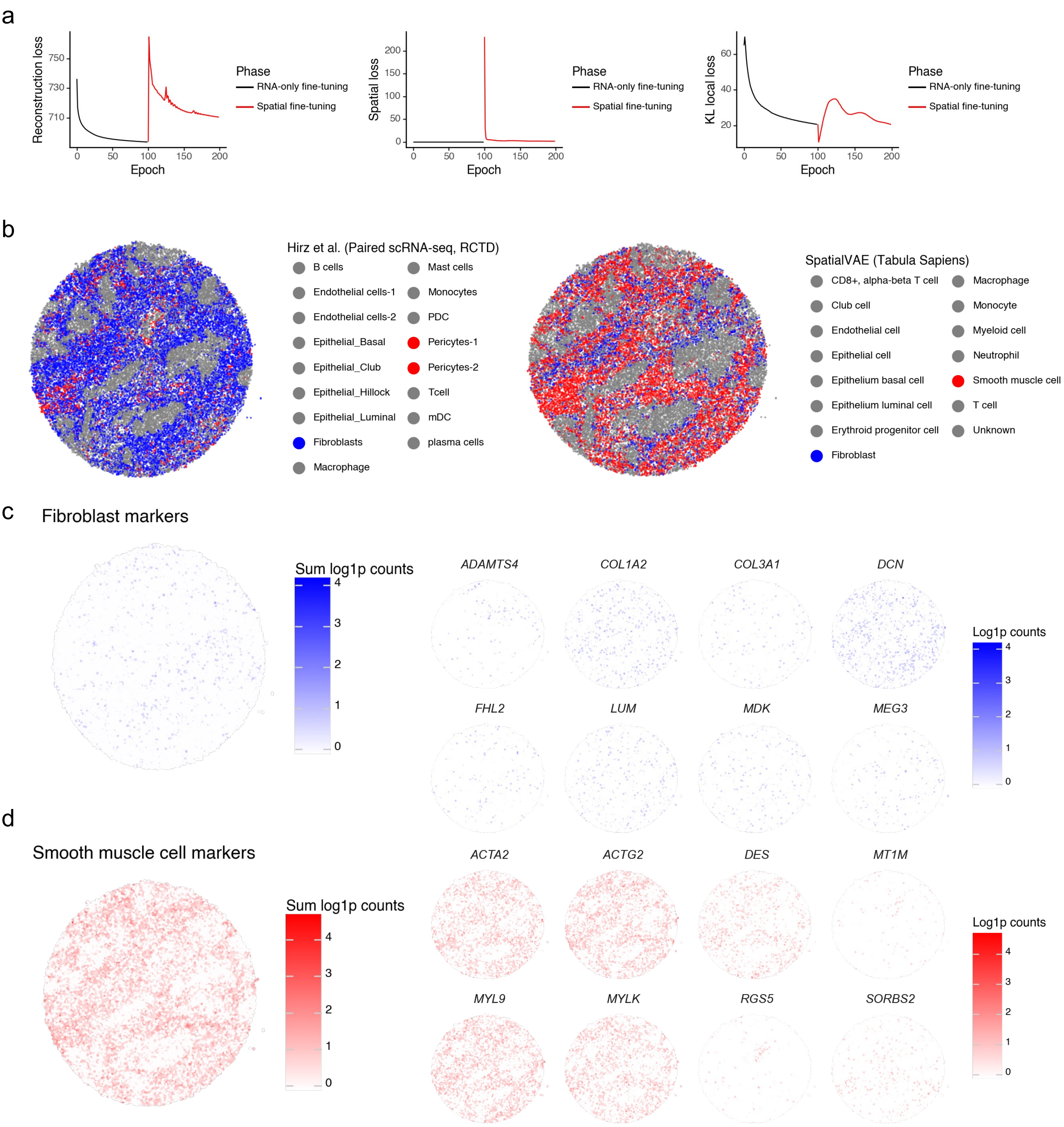


**Supplementary Figure 27: Spatially aware joint embedding of single-cell and spatial transcriptomics data of human prostate, related to Figure 6.**

(a) Visualizations of the training loss of the prostate SpatialVAE model. The overall loss objective is the sum of reconstruction loss (left), spatial loss (middle, zero for RNA-only models), and the KL local loss. Starting from the reference model, we first fine-tuned the model on the Slide-seqV2 data without the spatial loss until convergence (black curve, 100 epoch), then attached the spatial loss and fine-tuned with respect to the new objective (red curve). This is mainly to highlight the tradeoff between reconstruction accuracy and spatial consistency. Skipping the RNA-only fine-tuning step will not affect the performance of the final spatial model. (b-d) Mislabeling of the stromal populations in the original publication (Hirz et al.). (b) Hirz et al. and the Tabula Sapiens use slightly different cell type nomenclatures, where the ACTA2+ population is referred to as pericytes in Hirz et al. and as smooth muscle cells in the Tabula Sapiens. (c) Expression of fibroblast (DCN+, ACTA2- in Hirz et al.) marker genes. (d) Expression of smooth muscle cell (pericytes in Hirz et al., DCN-, ACTA2+) marker genes.
